# A human-specific modifier of cortical circuit connectivity and function improves behavioral performance

**DOI:** 10.1101/852970

**Authors:** Ewoud R.E. Schmidt, Hanzhi T. Zhao, Jung M. Park, Jacob B. Dahan, Chris C. Rodgers, Elizabeth M. C. Hillman, Randy M. Bruno, Franck Polleux

**Affiliations:** Department of Neuroscience, Columbia University, New York, NY 10032, USA; Department of Biomedical Engineering and Radiology, Columbia University, New York, NY 10032, USA; Mortimer B. Zuckerman Mind Brain Behavior Institute, Columbia University, New York, NY 10032, USA; Kavli Institute for Brain Science, Columbia University, New York, NY 10032, USA

**Author notes:** Address correspondence to: Franck Polleux, Ph.D., Columbia University, Department of Neuroscience, Mortimer B. Zuckerman Mind Brain Behavior Institute, Kavli Institute for Brain Science, Jerome L. Greene Science Center, 3227 Broadway, L5-050, MC 9853, New York, NY 10027.

## Abstract

The remarkable cognitive abilities characterizing humans are thought to emerge from our unique features of cortical circuit architecture, including increased feedforward and feedback connectivity. However, our understanding of the evolutionary origin and nature of these changes in circuit connectivity, and how they impact cortical circuit function and behavior is currently lacking. Here, we demonstrate that expression of the human-specific gene duplication *SRGAP2C* leads to a specific increase in feedforward and feedback cortico-cortical connectivity. Moreover, humanized SRGAP2C mice display improved cortical sensory coding, and an enhanced ability to learn a cortex-dependent sensory discrimination task. Our results identify a novel substrate for human brain evolution whereby the emergence of *SRGAP2C* led to increased feedforward and feedback cortico-cortical connectivity, improved cortical sensory processing and enhanced behavioral performance.

## INTRODUCTION

Recent studies have led to the identification of some of the genomic mechanisms and corresponding cellular and molecular substrates underlying human brain evolution. For example, during human brain development, prolonged neurogenesis and the expansion of a specific class of neural progenitors, the outer radial glia (oRG), have been proposed to lead to higher neuronal production (Fiddes et al., 2018; Hansen et al., 2010; Suzuki et al., 2018). In addition to this tangential expansion of the cortex that led to an increased number of cortical areas, human brain evolution has critically relied on changes in neuronal connectivity (Benavides-Piccione et al., 2003; Buckner and Krienen, 2013; Elston et al., 2001; Mohan et al., 2015; Seeman et al., 2018), including a shift towards increased cortico-cortical connectivity mediated by an expansion of supragranular layer 2/3 (Krienen et al., 2016; Zeng et al., 2012). However, while increased connectivity is considered a critical trait underlying human cognition, we currently lack the conceptual and experimental framework to understand how human-specific traits of cortical connectivity have emerged, how they impacted cortical function, and how they ultimately improved behavioral performance.

In recent years, a growing number of human-specific genetic modifiers have been identified, such as human-specific gene duplications (HSGDs) (Fortna et al., 2004; Sudmant et al., 2010), that were proposed to act as important modifiers of brain development. The first experimental test of this idea came from studies of the HSGD affecting the ancestral gene Slit-Robo GTPase Activating Protein 2A (*SRGAP2A*). Duplication of SRGAP2A specifically in the human lineage led to the emergence of the human-specific paralog *SRGAP2C* (Charrier et al., 2012; Fossati et al., 2016). When expressed in mouse cortical pyramidal neurons (PNs) *in vivo, SRGAP2C* inhibits the function of its ancestral copy *SRGAP2A*, resulting in changes in synaptic development mimicking those characterizing human cortical PNs. These include an increase in the density of both excitatory and inhibitory synapses received by layer 2/3 PNs, and neotenic features of excitatory and inhibitory synaptic development (Charrier et al., 2012; Fossati et al., 2016; Schmidt et al., 2019). These findings indicated that mouse cortical PNs humanized for SRGAP2C expression receive an increased number of synaptic inputs, similar to what is observed in humans (Benavides-Piccione et al., 2003; Elston et al., 2001). By targeting synaptic development, SRGAP2C may therefore act as a human-specific modifier of cortical connectivity in the human brain.

Here, we sought to investigate how SRGAP2C expression modifies cortical circuit structure and function by addressing a number of key outstanding questions: (1) what is the nature and the origin of the increased number of synaptic inputs? (2) how do these SRGAP2C-induced changes in synaptic development affect cortical connectivity, and (3) does SRGAP2C expression lead to functional changes of cortical circuits and subsequently behavior?

By employing a sparse, quantitative monosynaptic tracing method for layer 2/3 PNs, we found that humanization of SRGAP2C expression selectively increases feedforward and feedback cortical circuit connectivity. *In vivo* functional analysis of sensory processing then uncovered how SRGAP2C-induced changes in connectivity modifies neuronal response properties by increasing their response reliability and selectivity to sensory inputs. Finally, we show that mice humanized for SRGAP2C in forebrain excitatory neurons display an increased ability to learn a cortex-dependent texture discrimination task. We propose that SRGAP2C-induced changes in synaptic development represent a key substrate of human cortical circuit evolution that led to increased cortico-cortical connectivity, improved reliability of sensory coding, and enhanced learning in the context of sensory perception.

## RESULTS

### Quantitative mapping of neuronal connectivity in humanized SRGAP2C mice

To investigate how humanization of SRGAP2C expression modifies the structure and function of neuronal circuits, we developed a novel transgenic mouse line (*Rosa26-loxP-STOP-LoxP-SRGAP2C-HA* knockin) allowing for spatial and temporal control of SRGAP2C expression in a Cre-dependent manner (**Fig. 1, C and D, and fig. S1**). In this study, we will refer to SRGAP2C mice as animals that are heterozygous for the *Rosa26-SRGAP2C* allele, unless otherwise indicated.

**Fig. 1.**
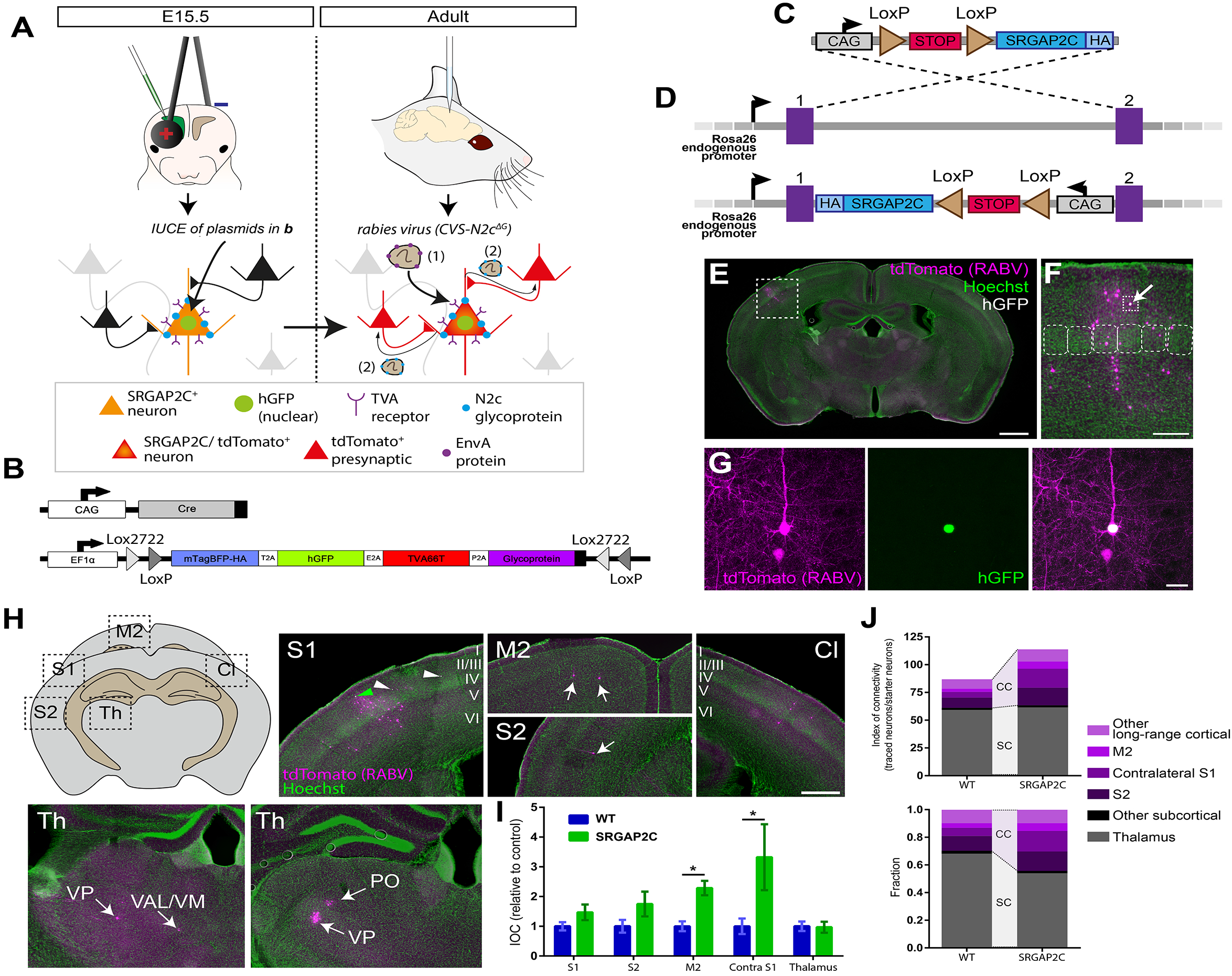
Sparse monosynaptic rabies tracing in humanized SRGAP2C mice. (**A**) The BHTG construct (see B), together with Cre recombinase, is targeted to layer 2/3 cortical pyramidal neurons in the primary sensory cortex (S1) by *in utero* electroporation (IUCE) at E15.5. When mice reach adulthood (>P65), stereotactic injection of RABV leads to initial infection of starter neurons expressing the BHTG construct (1), after which it spreads to presynaptically connected neurons (2). (**B**) BHTG and Cre constructs. (**C** and **D**) Design strategy for generating SRGAP2C conditional knock in mice. 3x HA tagged *SRGAP2C* was inserted into a *Rosa26* targeting vector (C), which contains a CAG promoter, a floxed STOP-Neomycin cassette, and *Rosa26* homology arms. Image not to scale. Using homologous recombination, the targeting vector was inserted between exon 1 and 2 of the *Rosa26* locus (D). (**E**) Coronal section stained for Hoechst (green) showing location of a starter neuron (indicated by dashed white box) in the barrel field of the primary sensory cortex (S1). Scale bar, 1 mm. (**F**) Higher magnification of dashed white box area in (E). Starter neuron (indicated by white arrow) is identified by expression of histone-GFP (hGFP, white). RABV infection is identified by expression of tdTomato (magenta). Rounded boxes indicate barrels in layer 4. Scale bar, 200 μm. (**G**) High magnification of starter neuron. Scale bar, 25 μm. (**H**) Anatomical location of RABV traced neurons. Boxes indicate brain regions where the majority of RABV traced neurons are located: the primary somatosensory cortex (S1) ipsilateral to the RABV injection site, secondary motor cortex (M2), secondary somatosensory cortex (S2), the primary somatosensory cortex contralateral to the injection site (Cl), and the thalamus (Th). For the thalamus, RABV traced neurons were located in the ventralanteriorlateral/medial (VAL/VM), ventral posterior (VP) and posterior (PO) subnuclei. Green arrowhead in S1 indicates RABV infected starter neuron. White arrowheads mark non-infected, electroporated neurons. White arrows mark RABV traced neurons. Roman numbers identify cortical layers. Scale bar, 500 μm. (**I**) Index of connectivity (IOC, number of traced neurons / number of starter neurons) for brain regions in (H), relative to control. (**J**) IOC (top) and fraction of inputs (bottom) for all RABV traced long-range inputs. Increased connectivity in SRGAP2C is selective for cortico-cortical (CC) inputs, but does not affect subcortical inputs (SC). Bar graphs plotted as mean ± s.e.m. \**P* < 0.05.

To determine the origin of increased connectivity received by layer 2/3 PNs following humanization of SRGAP2C expression during development, we developed a strategy using sparse *in utero* cortical electroporation (IUCE) combined with monosynaptic rabies tracing (Reardon et al., 2016; Wickersham et al., 2007) (**Fig. 1, A and B**). IUCE experiments were performed with a low amount of plasmid encoding Cre recombinase (see Methods), which led to a sparse population of layer 2/3 cortical PNs in the barrel field of the primary somatosensory cortex (barrel cortex) that express SRGAP2C in an otherwise wild-type brain. In addition, we co-electroporated a Cre-dependent plasmid expressing mTagBFP-HA, histone-tagged-GFP (hGFP), TVA^66T^ receptor, and N2c Glycoprotein (Miyamichi et al., 2013; Reardon et al., 2016) from a single cistron (**Fig. 1B**). TVA expression enables infection by EnvA pseudotyped G-deleted N2c rabies virus (CVS-N2c^ΔG^ [EnvA] RABV), while N2c glycoprotein facilitates trans-synaptic and retrograde spread of the virus (**Fig. 1A**). Hence, Cre-induced expression of SRGAP2C, hGFP, TVA and N2c marks a sparse population of SRGAP2C-expressing neurons and primes these ‘starter’ neurons for infection and trans-synaptic spread by RABV. When these animals reached adulthood (>P65), we performed stereotactic unilateral injection of RABV into the barrel cortex (**Fig. 1A**). One week after infection, robust expression of tdTomato was observed in neurons located in brain regions throughout the mouse brain (**Fig. 1, E to G**). A low number of layer 2/3 PNs expressing both tdTomato and nuclear hGFP were observed, indicating that initial infection by RABV was limited to a sparse number of starter neurons. Whole mouse brains were digitally reconstructed and the anatomical position of each traced neuron was mapped onto a reference atlas based on the Allen Institute Common Coordinate Framework (Allen Institute for Brain Science, 2015) (**Fig. S2A, and movie S1**). This allowed us to analyze each RABV traced brain within a common 3D reference space. Traced neurons were present in a large number of cortical and subcortical regions (**Fig. S2**, *n* = 10 for WT and *n* = 7 for SRGAP2C mice).

A large fraction of the traced neurons was observed locally, surrounding the starter neuron in the ipsilateral S1 (**Fig. 1H, and Fig. S2B**). We also consistently observed traced neurons in the ipsilateral secondary motor cortex (M2), ipsilateral secondary somatosensory cortex (S2), and the primary sensory cortex contralateral to the injection site. In addition to these cortical inputs, a sizeable number of inputs originated from subcortical regions. The majority of these neurons were located in thalamic nuclei, including the ventral-anterior lateral/medial (VAL/VM), ventral posterior (VP) and posterior (PO) nuclei (**Fig. 1, H and I, and fig. S4**), confirming previous studies showing that layer 2/3 PNs receive direct input from these thalamic nuclei, including inputs from VAL/VM (DeNardo et al., 2015; Viaene et al., 2010; Zhang et al., 2016). Together, these five brain regions (ipsilateral S1, S2, M2, contralateral S1 cortex, and ipsilateral thalamus) contained over 95% (95.8% ± 1.5% for WT, 96.3% ± 1% for SRGAP2C, mean ± s.e.m.) of all traced neurons.

### SRGAP2C selectively increases long-range cortico-cortical feedback connectivity

We next quantified the number of inputs received by layer 2/3 cortical PNs by calculating the Index of Connectivity (IOC), i.e. the number of traced neurons normalized by the number of starter neurons. Interestingly, when we assessed IOC for each brain region as a whole, we found that long-range cortico-cortical feedback projections from M2 and contralateral S1 were significantly increased in SRGAP2C mice compared to control wild-type mice (**Fig. 1I**, *P* = 2.54 × 10^−2^ for M2 and *P* = 1.56 × 10^−2^ for contralateral S1). This resulted in a specific increase in cortico-cortical inputs that SRGAP2C-expressing layer 2/3 PNs receive without a corresponding change in subcortical inputs (**Fig. 1J**). Connectivity originating in M2 was increased equally for both supra- and infragranular layers (*P* = 2.99 × 10^−2^ for supragranular and *P* = 4.1 × 10^−3^ for infragranular), maintaining the relative proportion between supra- and infragranular inputs (**Fig. 2, A to C**). In contrast, while for contralateral S1 we also observed an increase for both supra- and infragranular inputs (**Fig. 2, D to F**, *P* = 1.77 × 10^−2^ for supragranular and *P* = 1.39 × 10^−2^ for infragranular), the increase was relatively larger for inputs originating in the infragranular layers of contralateral S1 (**Fig. 2, E and F**). As a result, inputs from contralateral S1 onto *SRGAP2C*-expressing layer 2/3 PNs were more balanced between supra- and infragranular layer as opposed to WT neurons for which the majority of contralateral inputs originate in supragranular layers (**Fig. 2F**). Furthermore, when we analyzed the layer-specific contribution of S2, we found a significant increase for inputs originating in layer 4 of S2 (**Fig. 2, G to I**, *P* = 2.63 × 10^−2^), a projection that has recently been described as a non-canonical cortical feedback pathway between S2 and S1 (Minamisawa et al., 2018). Together, our data show that humanization of SRGAP2C expression specifically increases the number of long-range cortical feedback inputs layer 2/3 PNs receive, without affecting the number of subcortical inputs. Importantly, increased connectivity was not caused by differences in cortical depth or number of starter neurons (**Fig. S3B-D**). As expected, we observed a positive linear relationship between the number of infected starter neurons and the number of traced neurons. However, we found no correlation between IOC and number of starter neurons, which confirmed that our approach allows for the comparison of brains with small differences in the number of starter neurons (**Fig. S3C-D**).

**Fig. 2.**
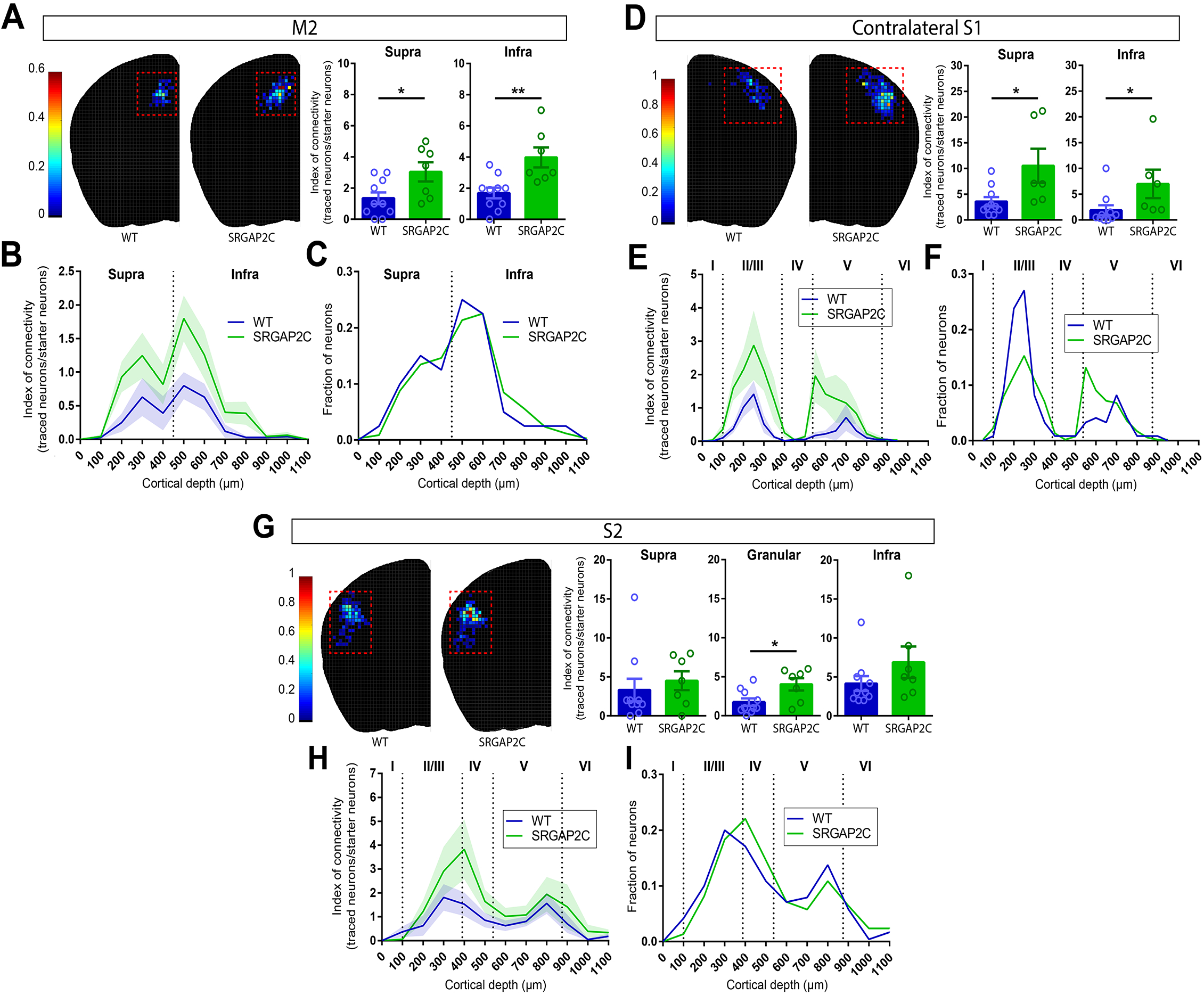
SRGAP2C expression modifies input distribution of long-range cortical inputs. (**A**) Index of connectivity (IOC, number of traced neurons / number of starter neurons) for supra- and infragranular layers in M2. Left: distribution of traced neurons in WT and SRGAP2C. Colors in density plot indicate IOC. Right: IOC for supra- and infragranular layers. (**B-C**) Distribution of traced neurons as a function or their cortical depth (**B**: IOC, **C**: fraction of total). (**D-I**) same as (A-C) for S1 contralateral to RABV injection site (D-F), and S2 (G-I). Bar graphs plotted as mean ± s.e.m. Open circles in bar graphs indicate data from individual mice. Shaded area indicates s.e.m. \**P* < 0.05, \*\**P* < 0.01.

### SRGAP2C increases local feedforward and feedback excitatory inputs

Besides long-range cortical and subcortical inputs, layer 2/3 cortical PNs in the barrel cortex receive a large number of inputs originating from within S1. These presynaptic neurons are located around the starter neurons (**Fig. 1H, and fig. 3, A and B**) and represent the majority of inputs layer 2/3 PNs receive (**Fig. 3C**). When quantifying IOC for S1 as a whole, we noticed a trend toward an increase in IOC (**Fig. 3B**, *P* = 0.109). Local inputs onto layer 2/3 PNs consists of excitatory inputs from other PNs and inhibitory inputs from interneurons of which the majority expresses either Parvalbumin (PV) or Somatostatin (SST). We wondered whether expression of SRGAP2C selectively changed the number of excitatory or inhibitory inputs layer 2/3 PNs receive since we previously demonstrated that SRGAP2C increased the density of both excitatory and inhibitory synapses found on apical dendrites of layer 2/3 PNs (Fossati et al., 2016). We performed post hoc immunofluorescent staining of all sections containing RABV traced neurons for PV and SST and quantified IOC for each of these subtypes (**Fig. 3, D and E**). Since PV and SST expressing interneurons provide the majority of inhibitory inputs onto layer 2/3 PNs, we classified RABV traced cells negative for these markers as excitatory (Tremblay et al., 2016). Interestingly, while the IOC or distribution of PV-positive and SST-positive cells was not different between WT and SRGAP2C expressing neurons (**Fig. 3E, and fig. S5, D to F**), the fraction of RABV-traced neurons provided by excitatory neurons was significantly increased in SRGAP2C-expressing compared to WT layer 2/3 PNs (**Fig. 3E**, *P* = 3.3 × 10^−2^).

**Fig. 3.**
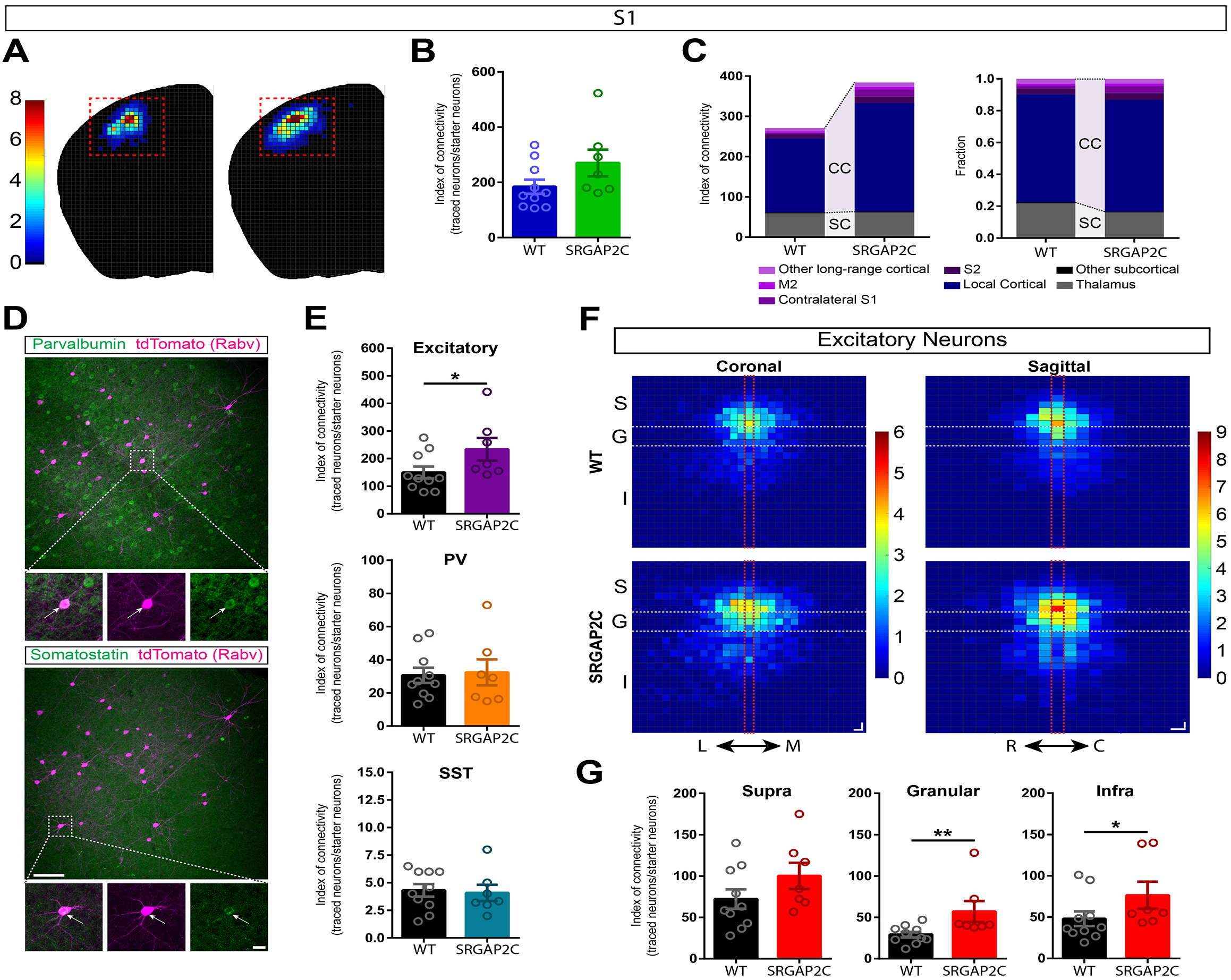
Expression of SRGAP2C leads to layer-specific increases in local excitatory connectivity. (**A**) Distribution of traced neurons in WT and SRGAP2C, colors indicate IOC. (**B**) IOC for primary somatosensory cortex (S1). (**C**) IOC (left) and fraction (right) for all RABV traced inputs, including local cortical inputs originating in S1. Increased connectivity in SRGAP2C is selective for cortico-cortical (CC) inputs but does not affect subcortical (SC) inputs. (**D**) Double immunohistochemistry of the same RABV traced brain section for Parvalbumin (top) and Somatostatin (bottom). Bottom panels represent higher magnification of area indicated by dashed boxes, showing co-labeling of Parvalbumin and Somatostatin with RABV-tdTomato. Scale bar large panel, 200 μm. Scale bar small panel, 20 μm. (**E**) IOC for excitatory, Parvalbumin (PV)- positive, and Somatostatin(SST)-positive RABV traced neurons in S1. (**F**) Density plots showing distribution of traced excitatory neurons relative to their closest starter neuron for coronal (left, L and M indicate lateral and medial orientation, respectively) and sagittal view (right, R and C indicate rostral and caudal orientation, respectively). Center bins aligned with relative position of starter neuron are indicated by red dashed lines. S, supragranular, G, granular, I, infragranular layers. For coronal, bin size = 50×50 μm. For sagittal, bin size = 50×100 μm. Colors in density plots indicate IOC. (**G**) IOC for excitatory neurons in supragranular, granular, and infragranular layers of S1. Open circles in bar graphs indicate individual mice. Bar graph plotted as mean ± s.e.m. \**P* < 0.05, \*\**P* < 0.01.

We next examined the laminar distribution of excitatory inputs. While we observed no differences for recurrent cortical inputs originating in the supragranular layers, we found a specific increase for local feedforward and feedback inputs originating in the granular and infragranular layers (**Fig 3, F and G, and fig. S5B**, *P* = 6.8 × 10^−3^ and *P* = 4.31 × 10^−2^ for excitatory neurons in the granular and infragranular layer, respectively). However, because the increase was larger for layer 4, this layer-specific increase altered the relative contribution of inputs from each layer, with a relative increase in layer 4 inputs and a relative reduction in inputs from layer 2/3 and upper layer 5 (**Fig. S5C**). The spatial distribution of locally traced neurons was not changed, indicating that the increased inputs received by SRGAP2C-expressing layer 2/3 PNs originated from the same local cluster of neurons and did not originate from more distantly located neurons in the antero-posterior or medio-lateral axes (**Fig. 3F and fig. S5A**). Together, these results show that SRGAP2C expression specifically increases the number of local excitatory feedforward and feedback inputs layer 2/3 PNs receive, without affecting recurrent cortical connectivity.

### SRGAP2C selectively increases synaptic density on apical dendritic domain

Connectivity in the mammalian neocortex is highly organized with neurons targeting specific layers of the cortex and selectively innervating neuronal subtypes and subcellular compartments (Douglas and Martin, 2004; Harris and Shepherd, 2015). The selective increase in cortico-cortical feedforward and feedback connectivity we observe in SRGAP2C-expressing layer 2/3 PNs may therefore arise from changes in synaptic density localized to specific dendritic domains. Our previous work focused on the analysis of apical oblique dendrites, but did not examine whether SRGAP2C expression affects synaptic development in other dendritic compartments (Charrier et al., 2012; Fossati et al., 2016). To extend these findings in our newly developed SRGAP2C mouse line, we used the same IUCE approach used for monosynaptic rabies tracing by electroporating a Cre-dependent plasmid encoding mTagBFP-HA (**Fig. 1B**) together with low amounts of Cre recombinase encoding plasmid, in heterozygous SRGAP2C or wild-type (WT) control mice. Electroporating low levels of Cre led to sparse labeling of layer 2/3 PNs in the barrel cortex (**Fig. S6, A and B**), allowing reliable quantification of dendritic spine density along distal tuft, apical oblique, and basal dendritic segments in optically-isolated neurons. While spine density was increased in the apical oblique and distal tuft segment of adult *SRGAP2C*-expressing neurons, no such increase was observed for basal dendrites (**Fig. S6C**, *P* = 1.92 × 10^−2^ for distal, *P* = 1.5 × 10^−3^ for apical oblique, *P* = 0.3 for basal). Spine size was similar between WT and SRGAP2C animals, showing that synapses are mature in SRGAP2C-expressing neurons after P65 (**Fig. S6D**) as previously reported (Charrier et al., 2012; Fossati et al., 2016).

### SRGAP2C increases neuronal response reliability and selectivity to sensory input

Local and long-range cortico-cortical connections play critical roles in the hierarchical processing of sensory information and response properties of layer 2/3 PNs (Adesnik and Naka, 2018; Petersen and Crochet, 2013). Selectively increasing feedforward and feedback cortico-cortical connectivity may therefore directly modify how layer 2/3 PNs process sensory information. To test this, we performed *in vivo* 2-photon Ca^2+^ imaging in WT and SRGAP2C mice crossed with Thy1-GCaMP6f and Nex^Cre^ mice (*n* = 4 WT and *n* = 3 SRGAP2C mice). Since the Nex^Cre^ mouse line induces recombination post-mitotically in all cortical pyramidal neurons (but not in interneurons or any non-neuronal cell types; (Goebbels et al., 2006)), this approach enabled us to image neuronal activity in layer 2/3 PNs (*n* = 962 neurons for WT and *n* = 618 for SRGAP2C) while ensuring that all imaged PNs express SRGAP2C.

Whisker stimulation (see Methods and **Fig. S7A**) was performed on awake mice for 5 seconds per trial, and repeated 24 times separated by 25 seconds inter-trial interval per imaging session (**Fig. 4, A to C**). This led to neuronal responses that were either time-locked to the onset of the stimulus (ON), responses that occurred during progression of the stimulus, or responses time-locked to the offset of the stimulus (OFF) (**Fig. 4, D and E**). Transients that were not time-locked to the onset or offset of the stimulus but occurred during the 5-second stimulus window often had longer response durations compared to ON or OFF responses (**Fig. 4D**). We refer to these as Sustained responses.

**Fig. 4.**
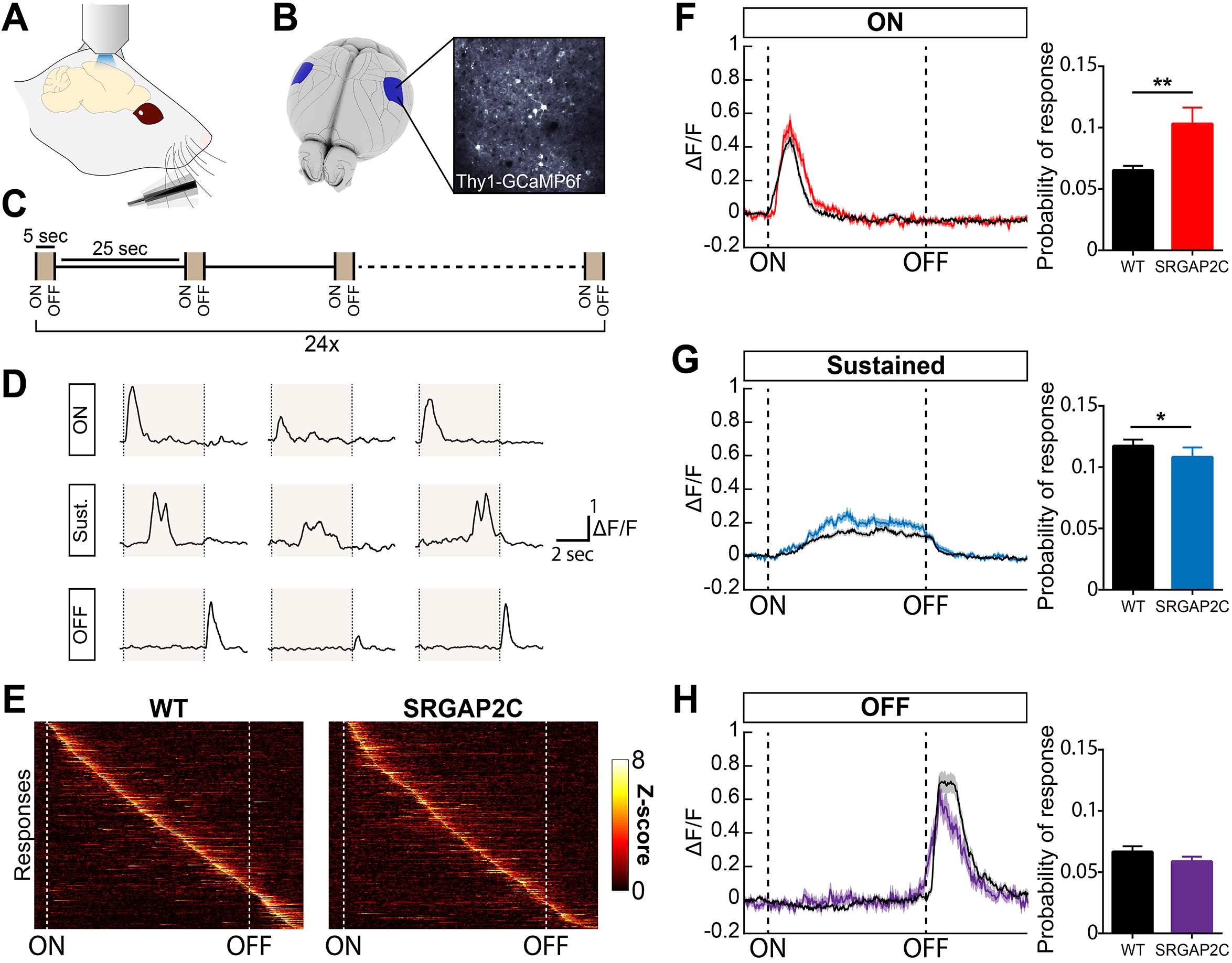
Humanization of SRGAP2C expression increases probability of neuronal responses to sensory stimulation. (**A** to **C**) Schema of experimental approach. (**D**) Singe-trial example responses. Shaded area indicates whisker stimulation. (**E**) Single-trial responses of responding layer 2/3 PNs converted to Z-scores and aligned to peak activity. (**F** to **H**) Left panels: average calcium traces (ΔF/F) for responses time-locked to the onset of the whisker stimulus (ON), during progression of the 5 sec stimulus (Sustained), or time-locked to the offset of stimulus (OFF). Shaded area indicates s.e.m. Right panels: response probability expressed as fraction of stimuli leading to a response. Bar graph plotted as mean ± s.e.m. \**P* < 0.05, \*\**P* < 0.01.

Overall, we observed no difference between WT and SRGAP2C mice regarding the fraction of layer 2/3 PNs that responded to the stimulus either as ON, OFF or Sustained responder neurons (**Fig. S7B**). We next measured the response probability for each neuron by calculating the fraction of trials that led to a response (ON or OFF), excluding neurons for which no responses were observed for any trials. Similar to previous studies (Clancy et al., 2015; Ramirez et al., 2014; Sato et al., 2007), overall sensory-evoked response probabilities were low in WT layer 2/3 PNs, with a long-tail distribution containing a small group of neurons that responded with high probability (**Fig. S7C**).

Interestingly, SRGAP2C expressing neurons had a significantly higher response probability specifically for responses time-locked to the onset of the stimulus, while a small but significant reduction in response probability was observed for Sustained responses and no significant changes in the probability of OFF responses (**Fig. 4, F to H**, *P* = 6.4 × 10^−3^ for ON, *P* = 4.31 × 10^−2^ for Sustained, and *P* = 3.21 × 10^−1^ for OFF). In addition, the duration of Sustained responses was significantly longer in SRGAP2C mice (*P* < 1 × 10^−4^), with a number of responses spanning nearly the entire stimulus (**Fig. 5, A to C**). Each whisker stimulus was separated by an inter-trial interval (ITI) of 25 seconds. Activity during the ITI was significantly reduced for SRGAP2C neurons for both the number of transients and transient amplitude (**Fig. 5, D and E**, median number of transients: 1.25 for WT and 0.5 for SRGAP2C, median transient amplitude: 1.26 for WT and 1.05 for SRGAP2C), which was not explained by differences in behavioral activity, such as whisking or grooming (**Fig. S7D**). Together with a higher response probability for the onset of the stimulus, this led to activity patterns that were more restricted to sensory stimulus epochs, resulting in an increase in response selectivity of SRGAP2C expressing layer 2/3 PNs (**Fig. 5D**).

**Fig. 5.**
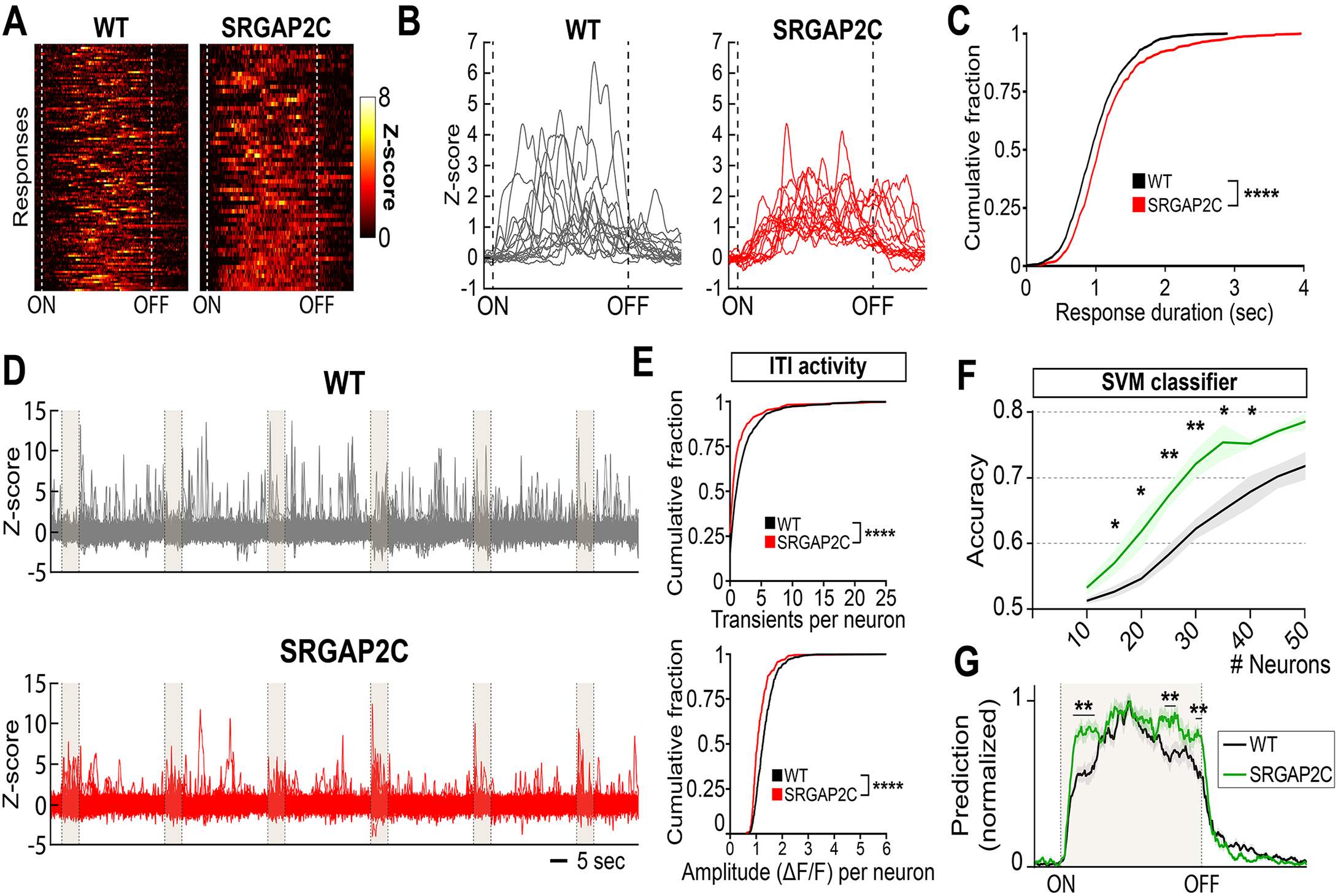
Humanization of SRGAP2C expression modifies neuronal response properties and accuracy of sensory coding. (**A**) Ten percent of single-trial Sustained responses with longest sustained activity converted to Z-scores and sorted by duration of response. ON and OFF dashed lines indicate stimulus onset and offset, respectively. (**B**) Bottom 15 responses shown in (A). (**C**) Cumulative probability distribution of Sustained response durations (time that z-score was greater than 1). (**D**) Z-scored example traces showing activity during six consecutive stimuli (shaded) and corresponding ITIs. (**E**) Cumulative probability distribution of Ca^2+^ transient number and amplitude during intertrial interval (ITI). (**F**) Support vector machine (SVM) accuracy in classifying presence or absence of whisker stimulus. Shaded area indicates s.e.m. (**G**) Normalized SVM prediction accuracy across time from stimulus ON to stimulus OFF for 25 neurons per field of view. Shaded area indicates stimulus time. \**P* < 0.05, \*\**P* < 0.01, \*\*\*\**P* < 0.0001.

Increased probability and selectivity of sensory-evoked neuronal responses suggests that SRGAP2C-expressing neurons encode sensory inputs more reliably. To determine whether SRGAP2C expressing neurons more accurately encode the presence or absence of a whisker stimulus, we trained a linear support vector machine (SVM) to classify for each time point whether the stimulus was ON or OFF. We analyzed accuracy of the classifier for a randomly selected group of neurons per field of view (FOV). This random selection was repeated 50 times after which we calculated average accuracy across these 50 iterations. We started with a random selection of 10 neurons per FOV and progressively increased this by including an additional 5 neurons. As expected, increasing the number of neurons used for training the SVM classifier increased its accuracy (**Fig. 5F**). Importantly, accuracy of the classifier was significantly higher for SRGAP2C neurons. When we examined the weights from the classifier grouped by ON, Sustained, and OFF responses, we found a significant increase in weights assigned to ON and Sustained responses (**Fig. S7E**). We next determined performance of the SVM classifier across time. Since the classifier is binary, it predicts for every time point whether the stimulus is ON or OFF. We aligned the time courses for all 24 stimuli and assessed the average accuracy for each time point across all 24 stimuli. We then normalized the results which allowed us to compare the relative prediction accuracy across time for WT and SRGAP2C expressing neuronal ensembles in each FOVs. For WT, the classifier gradually increased prediction accuracy and reached its maximum approximately halfway into the stimulus, after which it decreased again towards the offset of the stimulus. In contrast, for SRGAP2C-expressing neurons, the classifier was more accurate across the entire duration of the stimulus, with a significant increase in predicting stimulus onset as well more accurately predicting continuation of the stimulus until stimulus offset (**Fig. 5G**). Together, these results show that SRGAP2C expression in layer 2/3 PNs improves the accuracy of sensory coding by increasing response reliability and selectivity to whisker inputs.

### Whisker-based texture discrimination in humanized SRGAP2C mice

Long-range cortico-cortical projections play critical roles in motor learning and sensory perception (Gilad et al., 2018; Makino et al., 2017; Manita et al., 2015). In addition, response reliability and timing of cortical PNs play an important role in sensory processing and may represent a potential mechanism through which the cortex modifies its responsiveness to sensory inputs (Benedetti et al., 2009; Zuo et al., 2015). Increased feedforward and feedback cortico-cortical connectivity together with improved sensory coding accuracy may therefore directly modify, and potentially enhance, sensory processing in SRGAP2C mice. To investigate how this might affect behavior, we tested the ability of WT and Nex^Cre^-SRGAP2C mice to discriminate different textures in a whisker-based texture discrimination paradigm. Whisker-based texture discrimination relies on cortical processing in the barrel field of the somatosensory cortex, and requires long-range cortico-cortical projections from M2 to S1 (Guic-Robles et al., 1992; Manita et al., 2015). To allow for a potential increase in performance or learning ability, we increased the difficulty of the task by presenting mice with two different rough textures (see Methods), instead of more easily learned smooth versus rough texture discrimination (Park et al., 2020).

Water-restricted mice were trained in the dark to perform a two-alternative forced choice (2AFC) whisker-based texture discrimination task (**Fig. 6A-B, and movie S2**). Mice were randomly presented with rough (R200; grooves 200 μm apart) or less rough (R2000; grooves 2000 μm apart) textures at less than 1 cm from their whisker pad and, relying on only their whiskers, learned to respond by licking the left or right lick port, respectively, to obtain a water reward. Incorrect responses were punished with a timeout. Performance improved over sessions for both WT and SRGAP2C mice (*n* = 20 mice for WT and *n* = 18 for SRGAP2C mice, pooled from two independent cohorts). Following the last session, all whiskers facing the texture were trimmed which resulted in decreased, chance-level performance confirming that the mice from both groups performed the task in a purely whisker-dependent manner (**Fig. S8A**).

**Fig. 6.**
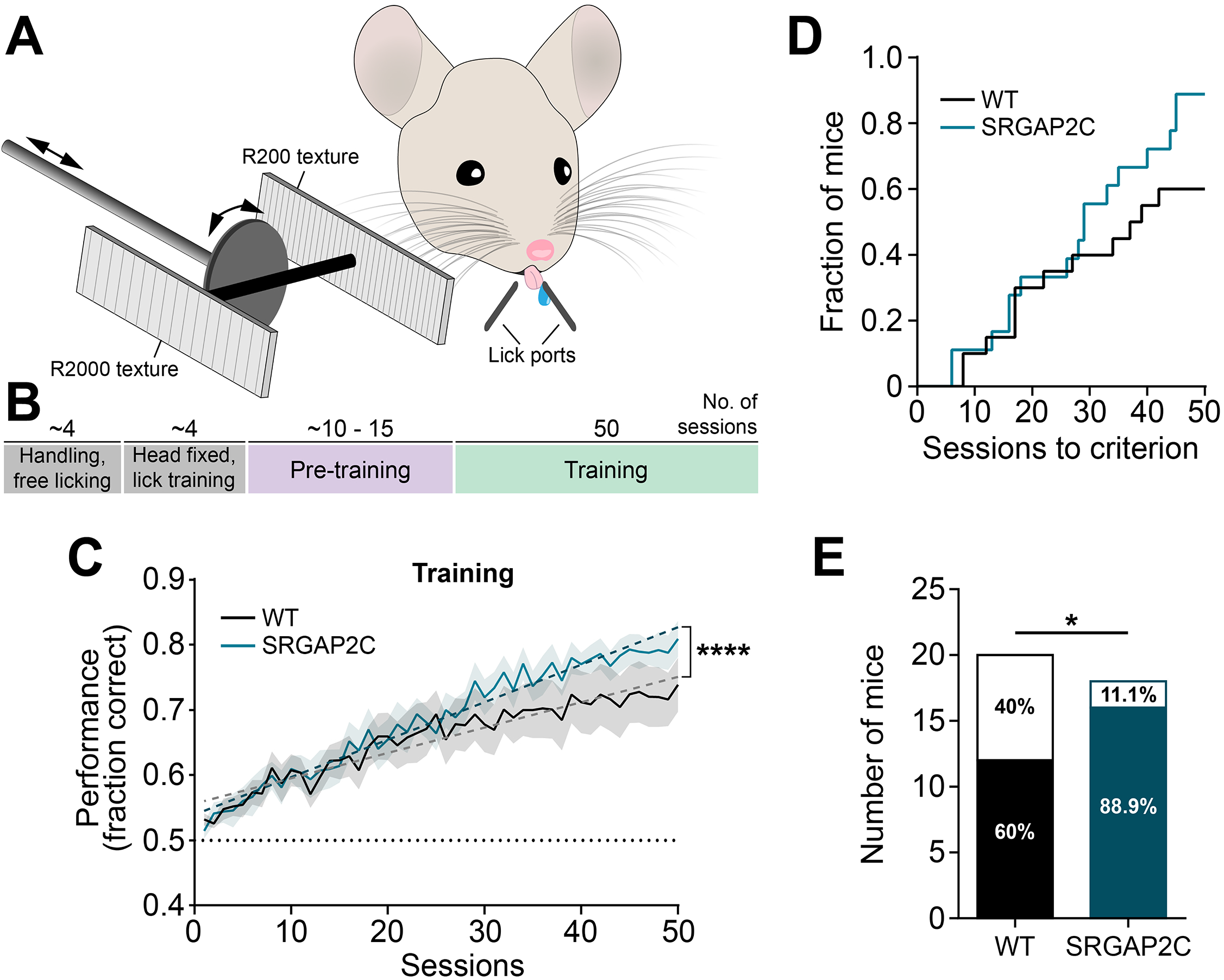
Humanized SRGAP2C mice display increased learning ability in texture discrimination task. (**A** and **B**) Schematic of whisker-based texture discrimination task. Distance between laser cut grooves for R200 and R2000 textures is 200 and 2000 μm, respectively. (**C**) Behavioral performance of WT and SRGAP2C mice shown as fraction of correct trials over sessions of training. Shaded area indicates standard error to the mean (s.e.m.). Linear regression indicated by dashed lines. Dotted horizontal line indicates chance level performance (50% correct). (**D**) Cumulative histogram for number of sessions to learning criterion (75% correct). (**E**) Number of learners (mice that reached 75% correct responses) and non-learners (mice that never reached 75% correct responses). Numbers in bar graphs indicate percentage of total number of mice tested. \**P* < 0.05, \*\*\*\**P* < 0.0001.

As a group, SRGAP2C-expressing mice displayed an increased learning rate, which was evident from a significant difference in slope of the performance curve (**Fig. 6C**, linear regression, slope WT = 0.0039, slope SRGAP2C = 0.0057, *P* < 0.0001). Due to the difficulty of the task, each group consisted of learners (>75% correct) and non-learners (**Fig. 6D**). SRGAP2C mice were significantly more likely to learn the task than WT mice (**Fig. 6E**, *P* = 4.34 × 10^−2^), an effect that was consistent for both independently tested cohorts (Cohort 1 learners: 62.5% for WT and 88.9% for SRGAP2C. Cohort 2 learners: 58.3% for WT and 90.9% for SRGAP2C). This result was neither explained by a difference in the number of sessions or trials the mice performed during the training stage, nor by a difference in number of sessions or trials run for pre-training, during which mice learn to alternate their licking between the left and right lick port when presented with the associated texture (**Fig. S8B**). However, we did find that SRGAP2C mice required significantly fewer sessions to reach criterion during pre-training (**Fig. S8C**, *P* = 1.35 x 10^−2^).

## DISCUSSION

The results presented in this study demonstrate that expression of the human-specific gene duplication SRGAP2C alters cortical circuit architecture by increasing the number of local and long-range cortical inputs onto layer 2/3 PNs, improves sensory coding by layer 2/3 PNs, and enhances sensory learning in a cortex-dependent texture discrimination task.

Our results reveal that changes in synaptic development regulated by human-specific SRGAP2C (Charrier et al., 2012; Fossati et al., 2016) cell-autonomously shapes synaptic connectivity within cortical circuits, and indicates that (1) the wiring principles governing connectivity in the mammalian neocortex are in part regulated by synapse availability on the postsynaptic side and (2) that changes in the relative contribution of cortico-cortical connectivity represents a substrate of the genomic process that shaped human brain evolution. It has previously been suggested that synapse availability is regulated by both synaptic density and spine neck length (Stepanyants et al., 2002), two features that are increased in mouse cortical PNs when SRGAP2C is expressed (Charrier et al., 2012; Fossati et al., 2016). Increasing these features specifically for excitatory inputs located on apical dendrites, which are the main recipients of cortico-cortical inputs (Shipp, 2007; Thomson and Bannister, 2003), is compatible with our main finding showing that humanization of SRGAP2C expression selectively increases local and long-range cortico-cortical connectivity received by layer 2/3 PNs.

In contrast to an increased number of presynaptic neurons providing excitatory inputs to layer 2/3 PNs, we observed no difference in the number of presynaptic PV or SST interneurons connected to layer 2/3 PNs expressing SRGAP2C. This is surprising in light of our previous work, which showed that excitatory and inhibitory synapse density is increased to a remarkably similar level upon humanization of SRGAP2C in layer 2/3 PNs (Fossati et al., 2016). Our current results may hint at two types of connectivity changes in SRGAP2C mice: an increase in the total number of excitatory inputs received by layer 2/3 PNs mediated by a larger number of presynaptic neurons accompanied by an increase in the number of inhibitory synaptic inputs originating from the same number of inhibitory neurons, potentially through increased axon branching by PV and SST interneurons. This model is supported by previous studies showing that, in response to changes in neuronal activity in excitatory neurons, interneurons regulate the number of inhibitory synaptic inputs through changes in axon branching (Chen et al., 2011; Pieraut et al., 2014).

By increasing cortico-cortical connectivity from deeper cortical layers as well as long-range feedback projections from more distant cortical regions (S2, M2, and contralateral S1), humanization of SRGAP2C expression selectively increases excitatory cortical feedforward and feedback inputs received by layer 2/3 PNs. While the precise function of many of these local and long-range cortico-cortical inputs remains partially understood, it has become clear that top-down feedback inputs can dynamically modify how cortical neurons respond to sensory stimulation by, among other features, changing the response reliability of primary sensory areas such as S1 (Larkum, 2013; Lee et al., 2008; Zagha et al., 2013). Furthermore, local feedforward inputs, such as from layer 4, play an important role in driving layer 2/3 PN activity in response to sensory input. Layer 4 is one of the main thalamo-recipient layers and relays sensory input to both superficial and deep layers of the cortex. However, while these inputs are numerous (between 300 to 400 layer 4 spiny neurons converge onto a single rat layer 2/3 PN (Lubke, 2003)), modeling the impact of these connections has suggested that the majority of layer 4 inputs would be unable to drive spike firing in layer 2/3 PNs, and that driving responses of layer 2/3 PNs relies on strong inputs from a small number of neurons (Lefort et al., 2009; Sarid et al., 2007). Increasing the number of layer 4 neurons that input onto layer 2/3 PNs may increase the probability that such strong inputs occur, and consequently increase the probability of layer 2/3 PN firing in response to whisker stimulation.

We did not observe increased local recurrent, or horizontal, connectivity among layer 2/3 PNs. Recruitment of local horizontal connectivity, while increasing activity in other layers, leads to inhibition within the layer itself, most likely through supra-linear recruitment of inhibitory neurons (Adesnik, 2018; Adesnik and Scanziani, 2010; Kapfer et al., 2007). Recruiting the same number of layer 2/3 PNs in response to whisker stimulation and maintaining similar horizontal connectivity could therefore provide further opportunity for feedforward and feedback cortical inputs to increase response reliability in the primary sensory cortex.

Together, results from these studies provide a potential explanation for how increased feedforward and feedback connectivity improves reliability of sensory coding by layer 2/3 PNs. However, additional mechanisms, such as altered synaptic plasticity, may provide potential alternative mechanisms for how SRGAP2C expression modifies neuronal response properties. For example, we observed reduced ITI activity of layer 2/3 PNs, making their responses more selective for sensory stimulation. Reduced ITI activity may be explained by increased inhibitory drive, although our previous work has shown that the magnitude of the increased synaptic density induced by humanization of SRGAP2C expression is remarkably similar for excitatory and inhibitory synapses, suggesting preservation of E/I balance (Fossati et al., 2016). However, our observation of increased response probability and sustained activity during sensory stimulation, together with reduced ITI activity, may suggest that the increase in excitatory synapse density in SRGAP2C-expressing layer 2/3 PNs leads to a simultaneous, perhaps homeostatic, reduction in synaptic strength. When excitatory drive is low, such as in the absence of a sensory stimulus, this would lead to a reduction in spontaneous firing rates. However, upon sensory stimulation, increased local and long-range cortical inputs are likely sufficient to overcome this reduction in synapse strength and therefore more reliably recruit layer 2/3 PNs upon sensory stimulation.

Expansion of the human neocortex, including a significant increase in cortico-cortical connectivity, is considered a critical trait underlying our remarkable cognitive abilities (Buckner and Krienen, 2013). Local and long-range cortico-cortical projections facilitate communication and integration of information across different functional areas of the neocortex. They play an essential role in cognition by regulating both the hierarchical processing of information, which is critical for transformation of sensory information and feature detection, as well as building predictive models of the external environment that enables the brain to make inferences about the nature of sensory inputs and relate these to appropriate behavioral responses (Adesnik and Naka, 2018; Gilbert and Li, 2013). As such, local and long-range cortico-cortical projections are involved for a multitude of behaviors, including the texture discrimination task employed in this study (Gilad et al., 2018; Makino et al., 2017; Manita et al., 2015). To perform this texture discrimination task correctly, mice have to learn to (1) discriminate two textures using their whiskers and (2) to learn an association between each texture and the correct lick port for reward. Strikingly, only 60% of wild-type mice learned this task (reaching performance of 75% correct choices) after 50 training sessions, illustrating that this behavior is cognitively demanding. Increased cortico-cortical connectivity and increased reliability of sensory coding may offer a likely explanation for this improvement in behavioral performance. More reliable sensory coding may allow for the detection of more subtle differences between textures. In turn, increased cortico-cortical connectivity, facilitating more effective communication between cortical regions such as M2 and S1, could subsequently allow for a more effective association of the texture with appropriate behavioral output.

In summary, our results identify a new substrate for human brain evolution: the emergence of SRGAP2C approximately 3 million years ago at the birth of the *Homo* lineage (Dennis et al., 2017), led to increased cortico-cortical feedback and feedforward connectivity, enhanced sensory coding and improved behavioral performance required to solve complex cognitive tasks that involve associative learning.

## Acknowledgements

The transgenic inducible knockin mouse model described here was developed in collaboration with genOway (Lyon, France). We thank Thomas Reardon and Samaher Fageiry for kindly providing RABV, Miyako Hirabayashi and Qiaolian Liu for excellent technical help, and Darcy Peterka and the Zuckerman Institute’s Cellular Imaging platform for instrument use and technical advice. We also thank members of the Polleux lab and Pierre Vanderhaeghen for valuable discussions and inputs.

## Author contributions

E.R.E.S. and F.P. conceived the experiments. E.R.E.S carried out RABV tracing and synaptic analysis, E.R.E.S. and H.T.Z. performed two-photon imaging experiments, C.C.R. and J.M.P. developed the texture discrimination setups, and E.R.E.S., J.P., and J.B.D. performed texture discrimination experiments. E.R.E.S, H.T.Z. and J.M.P. analyzed the data, R.B. advised on behavior experimental design, and E.M.C.H. and R.B. advised on data analysis. E.R.E.S. and F.P. wrote the manuscript.

## Data and materials availability

Reagents and other materials will be available upon request to F.P.

## Supplementary Materials

### Materials and Methods

#### Mice

All animals were handled according to protocols approved by the institutional animal care and use committee (IACUC) at Columbia University, New York. All mice used in experiments were adults (>P65), heterozygous for indicated transgenes, and were maintained on a 12h light/dark cycle. Nex^Cre^ (NeuroD6^tm1(cre)Kan^) mice (Goebbels et al., 2006) were obtained through Jax and induce recombination in dorsal telencephalic-derived postmitotic neurons giving rise to all pyramidal neurons throughout the cortex, hippocampus and amygdala but not in astrocytes, interneurons or microglial cells in these structures. Thy1-GCaMP6f mice (Dana et al., 2014) were obtained through Jax (C57BL/6J-Tg(Thy1-GCaMP6f)GP5.17Dkim/J) and stochastically express GCaMP6f in a subset of excitatory pyramidal neurons in various brain regions, including cortex and hippocampus.

Conditional SRGAP2C expressing mice were generated using homologous recombination in C57BL/6J mouse ES cells (see **Fig. S1** for details) in collaboration with genOway (France). A targeting vector containing a CAG promoter, *SRGAP2C-3xHA* cDNA, rabbit-globin poly-A, a LoxP-STOP-LoxP and neomycin selection cassette, and homology arms consisting of sequences between exon 1 and exon 2 of the *Rosa26* locus was constructed. In addition, Diptheria Toxin A (DTA) cDNA was placed downstream of the 3' homology arm for negative selection of non-recombined ES cell clones. Southern blot analysis was used to confirm homologous recombination of the targeting vector in ES cell clones. Recombined ES cell clones were injected into C57BL/6J blastocysts and re-implanted into OF1 pseudo-pregnant females to generate chimeric males. Male chimeras were subsequently bred with wild-type C57BL/6J females to generate an F1 population of SRGAP2C mice. Heterozygous mice (*Rosa26^SRGAP2C^*(F/+)) were confirmed carriers of the transgene by genomic PCR and Southern blot analysis (**Fig. S1**). Wild-type and SRGAP2C alleles are identified with PCR of genomic DNA using the following primers: 5’-CAATACCTTTCTGGGAGTTCT-3’ and 5’-CTGCATAAAACCCCAGATGAC-3’ for detection of the WT allele, and 5’-CATGGGGGATATGGCTTCC-3’ and 5’-GGAACATCGTATGGGTAAGCG-3’ for detecting the presence of *Rosa26* targeted SRGAP2C. For *in utero* electroporation experiments, mice were crossed once with the outbred strain 129S2/SvPasCrl mice (obtained from Charles River) to produce F1 hybrids females used to generate timed-pregnant females by crossing with *Rosa26^SRGAP2C^*(F/+) heterozygous males (on pure C57BL/6J). This strategy is used because pure C57BL/6J females often cannibalize their offspring at birth following *in utero* electroporation experiments but not these F1 (C57BL/6J; 129S2/SvPasCrl) females.

#### Western Blot

Cre-dependent expression of SRGAP2C-HA protein was analyzed by crossing SRGAP2C mice with Nex-Cre mice (Goebbels et al., 2006) to induce expression of SRGAP2C in all excitatory forebrain neurons. Cortical hemispheres were dissected and homogenized in ice-cold homogenization buffer (N-PER Neuronal Protein Extraction Reagent (ThermoFisher Scientific) with cOmplete Protease Inhibitor Cocktail (Roche), 10 μM MG-132 (Sigma-Aldrich), and Benzonase (EMD Millipore)) using a disposable Biomasher II (Kimble Chase). After homogenization, samples were incubated for 30 min at 4°C in homogenization buffer and subsequently centrifuged at 10,000 g for 30 min in a cooling centrifuge at 4°C. Samples were prepared in Laemmli buffer (Bio-Rad) containing 10% 2-Mercaptoethanol and boiled at 95°C for 5 min. Using SDS-PAGE, proteins were separated and then transferred to a polyvinylidene difluoride (PVDF) membrane (Immobilon-FL, EMD Millipore). Western blotting was performed using anti-HA primary (1:1000, Anti-HA.11, Biolegend) and anti-actin (1:5000, MAB1501, Millipore) together with goat-anti-mouse IgG conjugated to IRDye 800CW (1:10,000, Li-Cor) and goat-anti-mouse IgG conjugated to IRDye 680RD (1:40,000). Imaging of immunoblots was performed on an Odyssey CLx Imaging System (Li-Cor).

#### DNA constructs

The BHTG construct was generated by subcloning mTagBFP-3xHA, histone-GFP, TVA66T and the N2c glycoprotein together with 2A self-cleaving peptide sequences (see **Fig. 1B** for details) in between NheI and AscI cloning sites of the pAAV-Ef1a-DIO eNpHR 3.0-EYFP plasmid (Addgene plasmid # 26966) using Gibson assembly cloning. pCAG-Cre was generated in the Polleux lab, as previously described (Courchet et al., 2013).

#### *In utero* cortical electroporation

*In utero* cortical electroporation (IUCE) was performed at E15.5 on isoflurane anaesthetized timed-pregnant SRGAP2C or control female mice as previously described (Hand and Polleux, 2011), with the following modifications. Endotoxin-free DNA containing 1μg/μl of BHTG plasmid and 10-20 ng/μl Cre plasmid was injected into the ventricles of E15.5 embryos using a heat-pulled capillary. Electroporation was performed by applying 5 pulses of 42 V for 50 ms with 500 ms intervals using a 3 mm diameter platinum tweezer electrode (Nepa Gene) and a square wave electroporator (ECM 830, BTX). After placing embryos back into the abdominal cavity, the incision was closed using sutures and the mouse allowed to recover on a heating plate.

#### Virus injection

Adult mice were anaesthetized using isoflurane and placed in a stereotactic frame (Stoelting). A small burr hole was drilled over the barrel field of the primary sensory cortex (1.3 mm posterior and 3 mm lateral to Bregma (Paxinos and Franklin, 2001)) using a high-speed dental drill. A glass pipette (Drummond Scientific) was heat-pulled (Narishige PC-10) to produce a tip of approximately 10 μm in diameter. It was then filled with viral vector solution containing CVS-N2c^Δg^ [EnvA] RABV-tdTomato and lowered into the brain to a depth of 200-300 μm. Virus was subsequently injected at bouts of 25 nl at 2 nl/s with a 20s interval until a total volume of 400-500 nl was injected. The glass injection pipette was left for 2 min after the injection was completed after which it was slowly removed. The skin was closed using sutures and the mouse was allowed to recover on a heating plate. Mice were left for 7 days to allow enough time for RABV-tdTomato to reach sufficient levels of expression in both starter neurons and presynaptic inputs.

#### Sparse monosynaptic rabies tracing and whole brain reconstruction

##### Tissue preparation

After 7 days of RABV tracing, mice were anaesthetized with isoflurane and intracardiac perfusion was performed using 4% paraformaldehyde (Electron Microscopy Sciences) in PBS. Brains were isolated and incubated overnight in a 4% paraformaldehyde in PBS solution at 4°C. The following day, brains were washed in PBS and sectioned along the coronal plane at 100 μm using a vibrating microtome (Leica VT1200S). Approximately 100 sections were collecting spanning the most rostral part of the cortex to the cerebellum. Sections were stained using Hoechst 33258 (Sigma-Aldrich) and subsequently mounted on glass slides in Fluoromount-G aqueous mounting medium (ThermoFisher Scientific).

##### Image acquisition

Sections were first imaged on a Nikon SMZ18 stereo microscope with automatic stage using a SHR Plan Apo 1X at a zoom magnification of 4x. Stitching was performed directly after imaging using Nikon NIS-Elements software. Sections containing both tdTomato and hGFP positive cells were subsequently imaged by collecting Z-stacks on a Nikon A1 confocal microscope using a 10x Plan Apo NA 0.45 objective (Nikon) for identification of RABV starter neurons, which we identified by co-expression of hGFP and tdTomato. Additional confirmation of starter neurons was done by imaging these neurons again using a 40x Plan Apo NA 0.95 (Nikon) objective.

##### Section alignment and registration to atlas

Imaged sections were aligned using rigid body alignment in StackReg (ImageJ plugin) and when necessary manual adjustments were made using Adobe Photoshop. Neurons were subsequently counted and annotated using Cell Counter (ImageJ plugin) after which sections and coordinates were imported into 3ds Max (Autodesk). Landmarks were then manually placed at multiple anatomical locations (113 anatomical landmarks) corresponding to landmarks we assigned to a 3D mouse brain atlas imported from the Allen Brain Institute (Common Coordinate Framework 3). Landmarks were placed at anatomical regions easily identified in imaged sections, such as the first section containing hippocampus or the first section where the corpus callosum forms a continuous bundle. In addition, a single landmark indicating the point at which both cortical hemispheres meet was assigned to each section (designated as midline landmark). Using these assigned landmarks, together with custom-written scripts in MAXScript, the 1) section stack was resized to match the reference brain size, 2) based on the midline landmark each section was aligned along the dorsal/ventral axis of the reference brain, and 3) using anatomical landmarks each neuron was assigned to the corresponding location in the reference brain. Finally, in order to assign proper cortical depth to each neuron, the pial surface was traced and reconstructed for each section and the distance to the pial surface for each neuron was measured along the line that intersects the pial surface perpendicularly. Neurons were color coded according to their assigned brain region or their calculated cortical depth to manually confirm proper alignment to the reference brain. Finally, the 3D position and assigned brain region for each neuron was exported for subsequent analysis.

##### Identification of interneurons

Tissue sections were washed in PBS and subsequently blocked overnight at 4°C in blocking buffer (PBS containing 1% Triton X-100 (Sigma-Aldrich) and 5% goat serum (Gibco)). The following day, sections were washed 3x for 1 h in PBS containing 0.5% Triton-X100. Next, sections were incubated with primary antibody in PBS containing 0.5% Triton-X100 and 0.5% goat serum for 4 days at 4°C. Sections were then washed 3x 1 h in PBS containing 0.5% Triton-X100 after which they were incubated with secondary antibodies conjugated to Alex Fluor-488, 546, and 647 of appropriate species (1:500, ThermoFisher Scientific). After several washes in PBS sections were mounted on glass slides and mounted in Fluoromount-G. Z-stacks were collected by imaging on a Nikon A1 confocal microscope using a 10x Plan Apo NA 0.45 objective (Nikon) to identify neuronal identity for Rabv-traced neurons. Primary antibodies used were mouse anti-HA (1:1000, Anti-HA.11, Biolegend), rat anti-somatostatin (1:100, MAB354, Millipore), guinea pig anti-parvalbumin (1:200, 195004, Synaptic Systems), and rabbit anti-DsRed (1:500, 632496, Takara Bio).

##### Data analysis

Index of connectivity (IOC) was calculated for each animal by dividing the number of traced neurons by the number of starter neurons. Average IOC density maps showing the distribution of traced neurons across the brain were generated in MATLAB (Mathworks) by generating IOC density maps per animal and subsequently combining these maps to calculate an average IOC density map per genotype. For density maps showing distribution of neurons relative to their closest starter neuron, we determined, per animal, the closest starter neuron for each traced neuron in the primary sensory cortex (S1). We then calculated the relative medial-lateral and rostro-caudal position of each traced neuron with respect to the closest starter neuron. The majority of traced neurons in S1 form a dense cloud closely around the starter neuron. However, this approach may misclassify which starter neuron belongs to which traced neurons when multiple starter neurons are in close proximity of each other. Differences in the distance between starter neurons in a single brain for WT and SRGAP2C mice could therefore obscure a potential change in the distribution of traced neurons between WT and SRGAP2C mice. We therefore analyzed the minimum distance between starter neurons for brains with more than 1 starter neurons. We found no difference in the distribution of starter neurons between WT and SRGAP2C mice (distance from starter neuron to nearest other starter neuron: 300.4 μm ± 55.92 for WT and 357.1 μm ± 74.19 for SRGAP2C, mean ± sem). Furthermore, when we analyzed brains with only 1 starter neuron (*n* = 3 for WT and *n* = 2 SRGAP2C) we similarly did not observe a difference in the spatial distribution of traced neurons (data not shown).

#### Dendritic spine analysis

##### Dendritic spine quantification

Tissue sections were washed in PBS and subsequently blocked for 3 h at room temperature in blocking buffer (PBS containing 0.2% Triton X-100 (Sigma-Aldrich) and 5% goat serum (Gibco)). Next, sections were incubated with primary antibody in PBS containing 0.2% Triton-X100 and 5% goat serum overnight at 4°C. The next day, sections were washed 3x for 30 min in PBS containing 0.2% Triton-X100. Sections were then incubated for 1 h using with secondary antibodies conjugated to Alex Fluor-546 or 647 of appropriate species (1:500, ThermoFisher Scientific). After several washes in PBS sections were mounted on glass slides using Fluoromount-G. Imaging of dendritic spines was performed using a Nikon A1 confocal microscope. First, low magnification Z-stack images were collected using a 20x Plan Apo NA 0.75 objective (Nikon) to visualize the entire dendritic tree of optically isolated neurons. Next, appropriate terminal branches were selected for distal, apical oblique, and basal dendritic segments, for which we collected Z-stack images using a 100x H-TIRF, NA 1.49 objective (Nikon).

##### Data analysis

Nikon NIS-Elements was used to generate a z-stack maximum intensity projection of selected dendritic branches. We then quantified spine density and head size by tracing the dendritic segment to measure the dendritic length and drawing ROIs around the spine head as visualized by the mTagBFP-HA filler signal.

#### Two-photon calcium imaging

##### Surgery and image acquisition

Adult mice were anaesthetized using isoflurane and injected with buprenorphine (0.1 mg/kg body weight), after which the dorsal skull was exposed and cleaned with a razor blade. A craniotomy was performed and a glass window was placed over the barrel field of the primary sensory cortex. The window was glued in place using cyanoacrylate glue, after which a custom head plate was secured onto the skull using both cyanoacrylate glue and dental acrylic cement. Mice were allowed to recover for a minimum of 7 days, after which imaging in awake mice (*n* = 4 WT and *n* = 3 SRGAP2C) was conducted on a Bergamo II two-photon microscope (Thorlabs) running Scanimage, using a 16x 0.8 NA objective (Nikon) and 920-nm wavelength Ti-Sapphire laser (Coherent). Imaging was performed 150 – 250 μm below the pial surface at 30 Hz and 512 x 512 pixels covering 830 μm x 830 μm. For whisker stimulation trials a transparent acrylic rod was placed next to the right whisker pad at a fixed distance of 2 mm. Every stimulus trial consisted of a 10 seconds pre-stimulus, 5 second stimulus, and 15 second poststimulus period. Whisker stimuli were applied by vibrating the bar at 25Hz, a speed of more than 100 mm/s, and an amplitude of over 1 mm. To identify the responding region in the barrel field of primary sensory cortex contralateral to the stimulated whisker pad, we first performed a wide-field imaging experiment in which the entire region under the cranial window was imaged during multiple stimulus trials. The hemodynamic component was calculated and the neural signal was isolated, as previously described (Ma et al., 2016). Subsequent two-photon imaging was performed on 2 to 3 non-overlapping fields of view per mouse within the center of the activated area. Activity of the animal during imaging was monitored using an infrared camera.

##### Image processing and analysis

Non-rigid motion correction of acquired images was performed using NoRMCorre, as previously described (Pnevmatikakis and Giovannucci, 2017). Regions of interest (ROIs) were manually drawn over soma of individual neurons. All subsequent analyses were done using custom-written MATLAB code, which is available upon request. Time-courses were calculated as the mean of all pixels within the ROI. Neuropil signal was estimated by dilating neuronal ROIs by 4 pixels to form a ring-shaped neuropil ROI around the soma of each neuron. To remove neuropil contamination, we subtracted the ΔF/F neuropil signal from the ΔF/F neuronal signal. To avoid subtraction of signal not considered neuropil contamination, we excluded pixels in the neuropil ROI that contained transients exceeding two times the standard deviation of the difference between the neuropil and neuronal signals, as previously described (Peters et al., 2014). To identify the response type per stimulus for each neuron, we aligned all time-courses to their respective stimulus onset and performed K-means clustering with 60 clusters and correlation as the distance metric. Common time-courses were identified by running K-means 200 times with random initialization. The resulting outputs of the repeated K-means were clustered a final time to obtain a set of representative responses. Clusters with responses where the onset time was time-locked with the start or end of the whisker stimulus were determined to be an on- or off-type response, respectively. Those occurring during the duration of stimulus were grouped as Sustained-type responses. Neuronal responses were subsequently assigned to a cluster by evaluating the Pearson’s correlation between the time-course and each basis time-course. The highest correlation value greater than 0.4 was determined to be the representative time-course. Duration of Sustained responses were calculated as the time where z-scored time-courses were greater than 1. To characterize activity during the inter-trial interval (ITI), transients were identified by finding the local maxima during ITI periods. Peaks were constrained to having widths of at least two frames, a minimum ΔF/F of 0.5, and a relative increase in ΔF/F of 6 times the standard deviation compared to the preceding local minimum. A frame-to-frame sliding window correlation analysis of webcam images was done to determine periods and duration of active behavior, such as whisking or grooming.

#### Support vector machine classification

Time courses for all neurons were z-scored, after which for each field of view (FOV) a random number of 10 neurons were selected to train a linear support vector machine (SVM) to classify between the presence (ON) and absence (OFF) of a whisker stimulus. For each FOV, the SVM was performed 50 times, each time with a new randomly selected group of 10 neurons. Each iteration was performed with 5-fold cross validation using 80% of time points for training and the remaining 20% for testing. Accuracy was calculated as the average across all 50 iterations. We repeated this procedure by progressively increasing the number of neurons per FOV used for the SVM by 5. In order to calculate accuracy of the SVM across time, we calculated the average accuracy for each time point across all 24 stimuli. This generated an average accuracy value for each time point of the time course. We grouped these results for WT and SRGAP2C FOVs to generate an average accuracy across genotypes and normalized the results to the maximum (set to 1) and minimum (set to 0) values. Weights, defined as the orthogonal vector to the separating hyperplane, were normalized per FOV to the maximum absolute weight, i.e. the absolute distance farthest away from the hyperplane. All SVM analyses were performed using custom written MATLAB code (Mathworks).

#### Texture-discrimination task

##### Surgeries

Adult mice were anaesthetized using isoflurane and injected with meloxicam, after which the dorsal skull was exposed and cleaned with a scalpel. A custom-designed steel head plate (Wilke Enginuity) was subsequently secured onto the skull using Metabond (Parkell).

##### Behavior setup

The behavioral apparatus was contained within a black box (Foremost) with a light-blocking door. A stepper motor (Pololu) rotated custom-designed textures in position after which they were advanced into the final stimulus position within approximately 1 cm of the right whisker pad of the mouse using a linear actuator (Actuonix L12-30-50-6-R). Textures were laser cut from acrylic sheets to a dimension of 16 x 33 mm. Vertical grooves of approximately 500 μm deep and 350 μm wide were laser cut into the acrylic textures at a spacing of 200 or 2000 μm apart. These were identified as R200 and R2000, respectively. Rewards (~5 μL of water) were delivered by opening a solenoid valve (The Lee Co. LFAA1209512H) that allowed water to flow from a reservoir to the lick port, which was made of a stainless steel tube (McMaster). Two lick ports were positioned in front and slightly to the left and right of the mouse's mouth, and licking was registered using capacitive touch sensors (Sparkfun MPR121). Between trials, a white light (LE LED 1800016-WW-US) was activated to prevent mice from fully dark-adapting. This prevented mice from using visual cues in performing the task. A computer fan (Cooler Master 80 mm Silent Fan) blew air over the texture away from the mouse, preventing the mouse from picking up olfactory cues. In addition, the textures were regularly cleaned with 70% ethanol. We never observed mice exploiting auditory or vibrational cues from the motors and thus no masking noises were necessary.

All aspects of the task were controlled by an Arduino Uno. A desktop PC chose the stimulus and correct response and logged all events read from the Arduino to disk using custom Python code. The training parameters for each mouse were stored in a custom-written Django database and updated manually or semi-manually by the experiments depending on each mouse’s progress.

##### Mouse training and testing

Experimenters were blind to the genotype of mice during every step of training and testing. Mice were denied water access in the home cage and learned to receive their water during behavioral training. For each mouse we closely monitored water intake, weight, and general health to ensure they did not suffer from dehydration. *Ad libitum* water was provided as necessary to prevent adverse effects on health. Mouse training was performed according to the following pipeline (also see **Fig. 6B**). 1) Handling and free licking. During this stage, mice were handled to become familiar with the experimenters. By placing mice in the set up without head-fixing, and allowing them to drink freely from the water pipes, they also became familiar with the behavioral set up. This stage required on average 4 sessions. 2) Head-fixed lick training. Mice were head-fixed directly in front of a single lick pipe and received a water reward for every lick. Next, mice were presented with two lick pipes on the left and right side of their mouth and learned to lick alternatively from each of them. This lick training started with mice alternating after every ten licks and was gradually decreased to a single lick on each side. This stage required on average 4 sessions. 3) Pre-training. Here, mice were presented with the complete trial structure for the first time, i.e. textures were presented and mice were only rewarded for correct responses. Incorrect responses were punished with a timeout. Presentation of the same texture was repeated until mice responded correctly, after which the other texture was presented. Thus, mice could perform at 100% by alternating responses from trial to trial. The timeout was initially 2 s and gradually increased to 9 s as mouse performance increased. This stage required between 10 – 15 sessions. 4) Training. Each session began with 45 trials of pretraining (as in step 3) to verify mice were able to lick both lick ports. After these initial pre-training trials, presentation of textures was randomized. The software continuously monitored mouse performance for biases, where mice stopped alternating between lick pipes and focused on one lick pipe only. When a strong bias was detected, presentation of the textures stopped being random and were presented to counteract the bias: if mice responded ≥20% more on one side than the other, the software would deliver only trials opposite to the biased side. If mice showed a significance perseverative bias (*P* < 0.05, ANOVA), the software would deliver alternating trials. For analysis, non-randomized trials were excluded and only randomized trials were used to assess performance.

Finally, to assess that mice relied on their whiskers to discriminate between texture, whiskers facing the texture side were fully trimmed. If performance did not fall to chance (<60% correct), mice were excluded from analysis. One mouse was excluded based on this criterion.

#### Statistics

Statistical analysis was performed using Prism 6 (Graphpad Software). Normality was checked using Kolmogorov-Smirnov test. A non-parametric test (Kruskal-Wallis with post-hoc Dunn's multiple comparison test or Mann-Whitney U test) was used when distribution deviated significantly from normality. A test was considered significant when *P* < 0.05. For RABV tracing, 10 WT and 7 *SRGAP2C* mice were obtained from a minimum of four independent litters. For *in vivo* spine analysis, data was obtained from at least six animals from a minimum of three independent litters. For two-photon microscopy experiments, 4 WT and 3 SRGAP2C mice were obtained from two independent litters. For the texture-discrimination task, 20 WT and 18 SRGAP2C mice were tested across two separate cohorts from a minimum of four independent litters. Significance was tested using a chi-squared test. Independent data points shown denote data from individual animals.

---

**Fig. S1.**
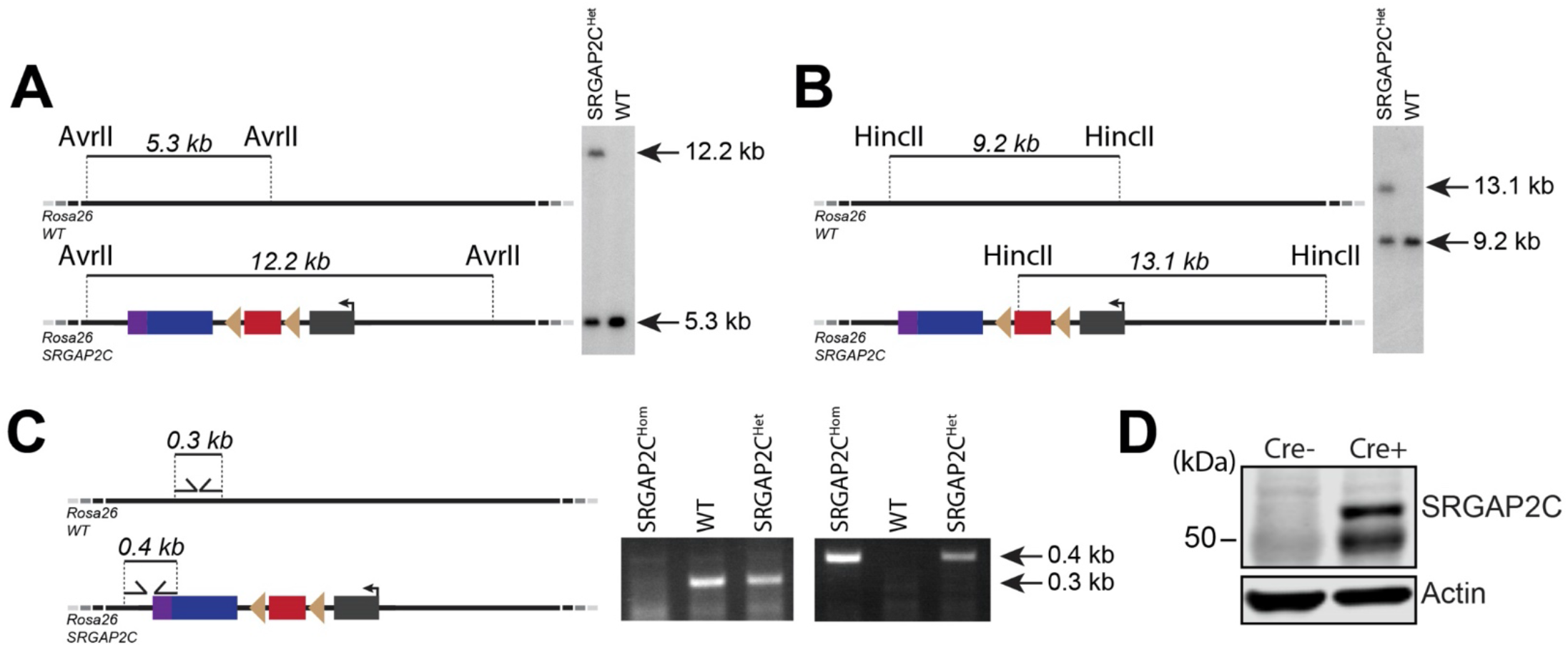
Generation of an inducible, humanized SRGAP2C transgenic mouse line. (**A** and **B**) Verification of SRGAP2C targeting in mouse embryonic stem cells using Southern blot analysis with probes that distinguish the targeted allele (12.2 kb in (A), 13.1 kb in (B)) from the wildtype allele (5.3 kb in (A), 9.2 kb in (B)). (**C**) Mice were genotyped by genomic PCR using the forward and reverse primers indicated that distinguish the WT Rosa26 allele or the SRGAP2C allele. (**D**) Western blot probed with anti-HA antibody of adult (P30) cortex isolated from SRGAP2C heterozygous conditional knock-in mice crossed with heterozygous Nex^Cre/+^ mice (Cre+) or wild-type littermate (Cre-). The presence of Cre induces SRGAP2C-HA expression. Without Cre, no SRGAP2C was detected. Anti-Actin antibody was used as loading control.

**Fig. S2.**
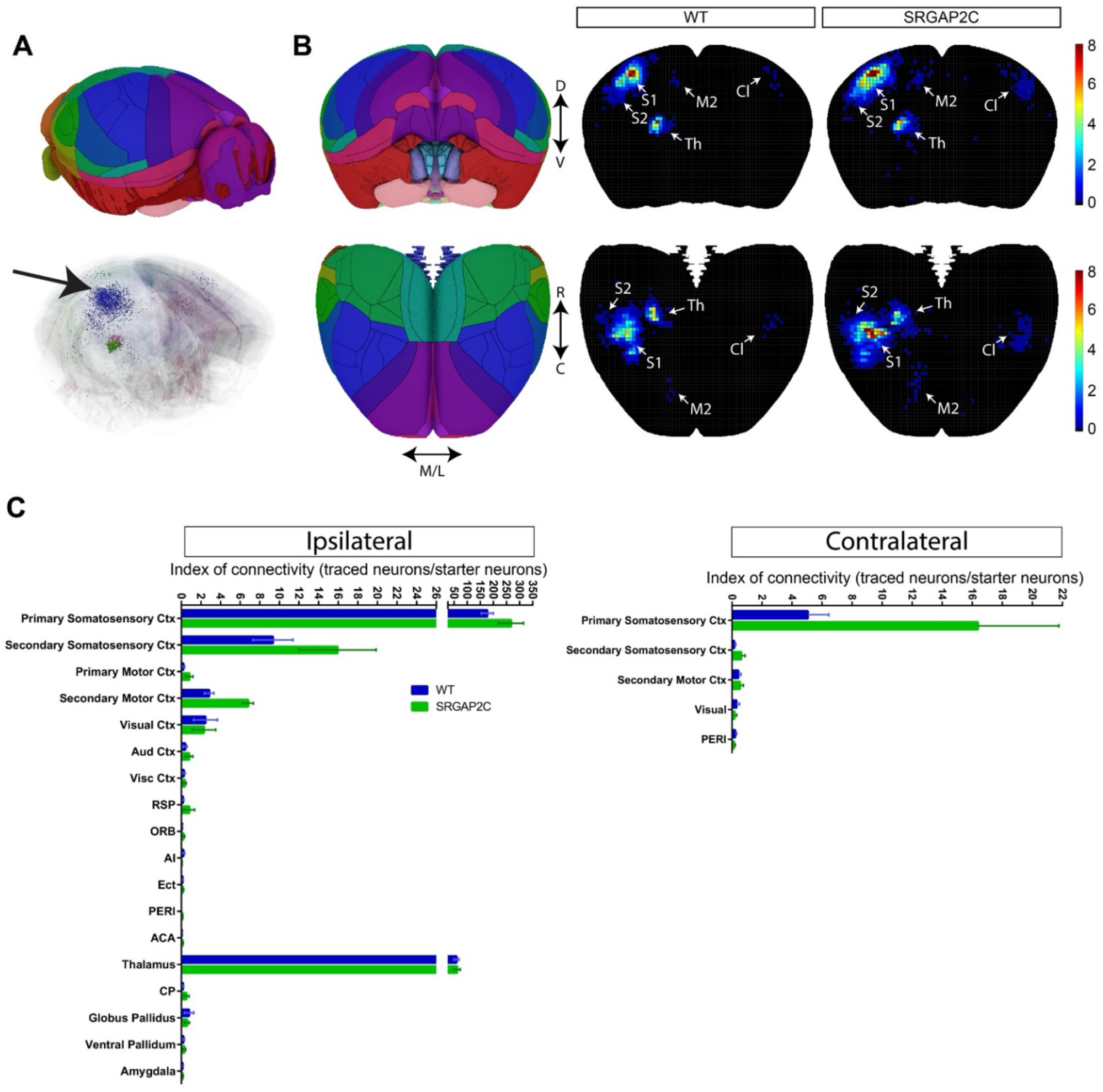
Brain regions containing RABV traced neurons. (**A**) Reference brain (top) based on Allen Reference Atlas. Digital reconstruction of RABV traced brain and registration onto reference brain. Black arrow indicates location of starter neurons in barrel field of S1. (**B**) Density plots showing distribution of traced neurons in WT and SRGAP2C mice. Colors in density plot indicate index of connectivity (IOC): number of traced neurons / number of starter neurons). (**C**) IOC for brain regions ipsilateral and contralateral to the injection site. RSP, retrosplenial area, ORB, Orbital cortex, Ai, Agranular Insular cortex, Ect, Ectorhinal cortex, PERI, Perirhinal cortex, ACA, Anterior Cingulate cortex, CP, Caudate-putamen. Bar graphs plotted as mean ± s.e.m.

**Fig. S3.**
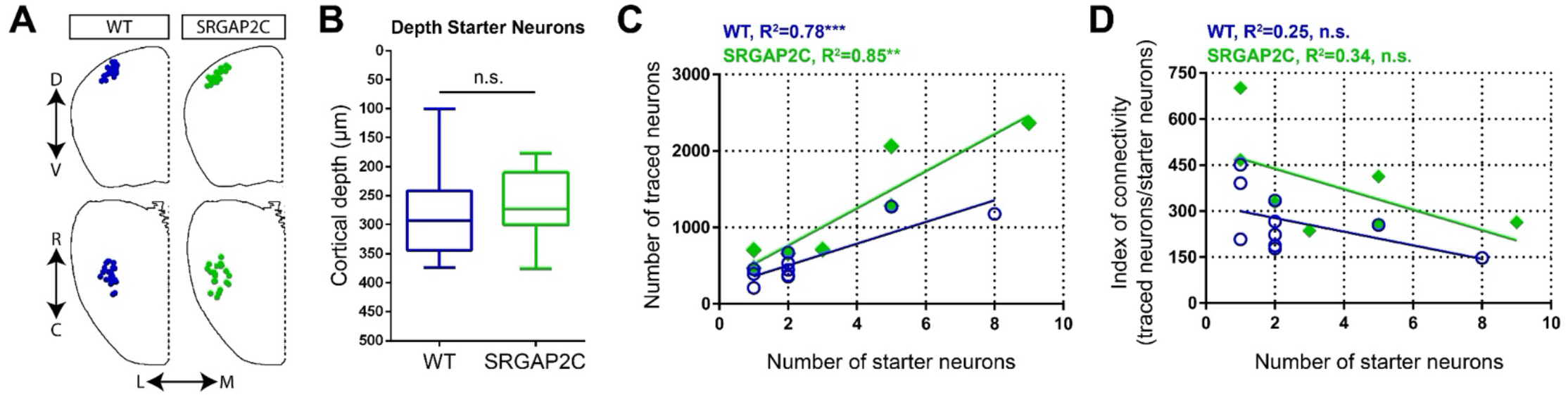
Connectivity changes are not caused by differences in cortical depth or number of starter neurons. (**A**) Location of starter neurons (*n* = 26 starter neurons, 10 mice for WT, and *n* = 26 starter neurons, 7 mice for SRGAP2C). (**B**) Cortical depth of starter neurons measured as distance from pial surface is not different between WT and SRGAP2C mice. Data shown as box-and-whisker plots. Center line indicates median, box edges represents first and third quartiles, and whiskers represent minimum and maximum values. (**C**) Correlation between number of RABV infected starter neurons and RABV traced neurons (Pearson's correlation coefficient *r* = 0.88, *P* = 7 × 10^−4^ for WT, and *r* = 0.92, *P* = 3.2 × 10^−3^ for SRGAP2C). (**D**) No correlation was observed between IOC and number of RABV infected starter neurons per brain (Pearson's correlation coefficient *r* = −0.5, *P* = 0.14 for WT, and *r* = −0.58, *P* = 0.17 for SRGAP2C).

**Fig. S4.**
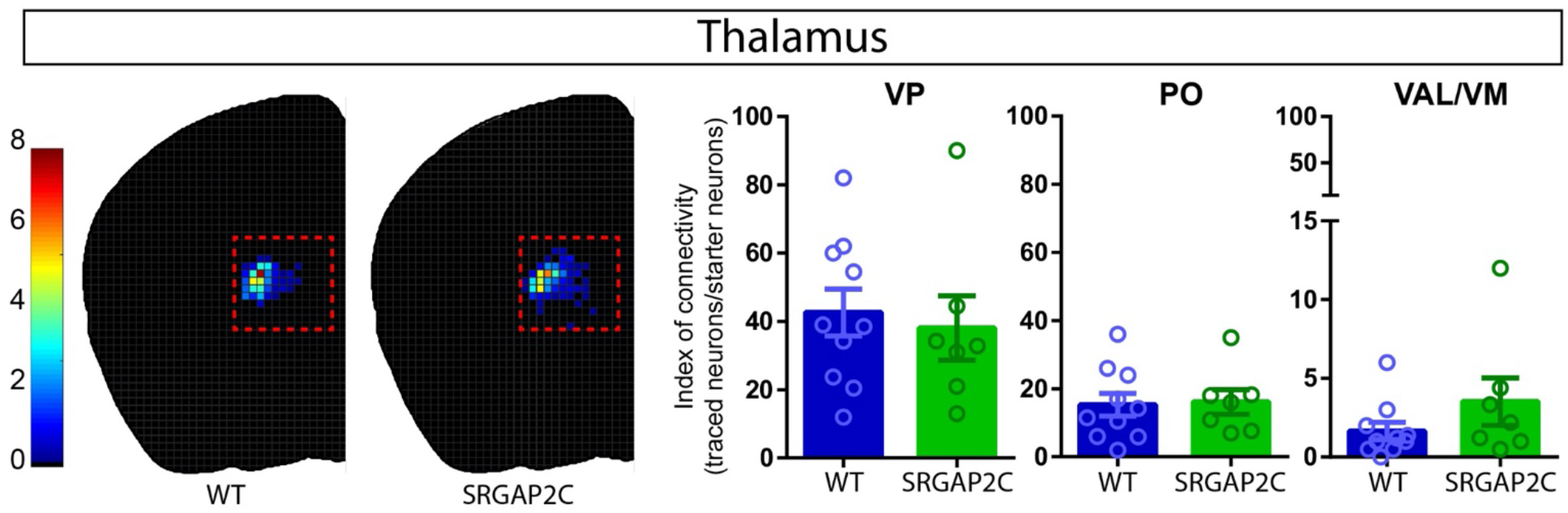
Distribution of RABV traced neurons in the thalamus. Index of connectivity (IOC, number of traced neurons / number of starter neurons) for traced neurons in the thalamus. Left: distribution of traced neurons in WT and SRGAP2C, colors indicate IOC. Right: IOC for Ventralanteriorlateral/medial (VAL/VM), Ventralposterior (VP), and Posterior (PO) thalamic subnuclei. Bar graphs plotted as mean ± s.e.m. Open circles in bar graphs indicate individual mice.

**Fig. S5.**
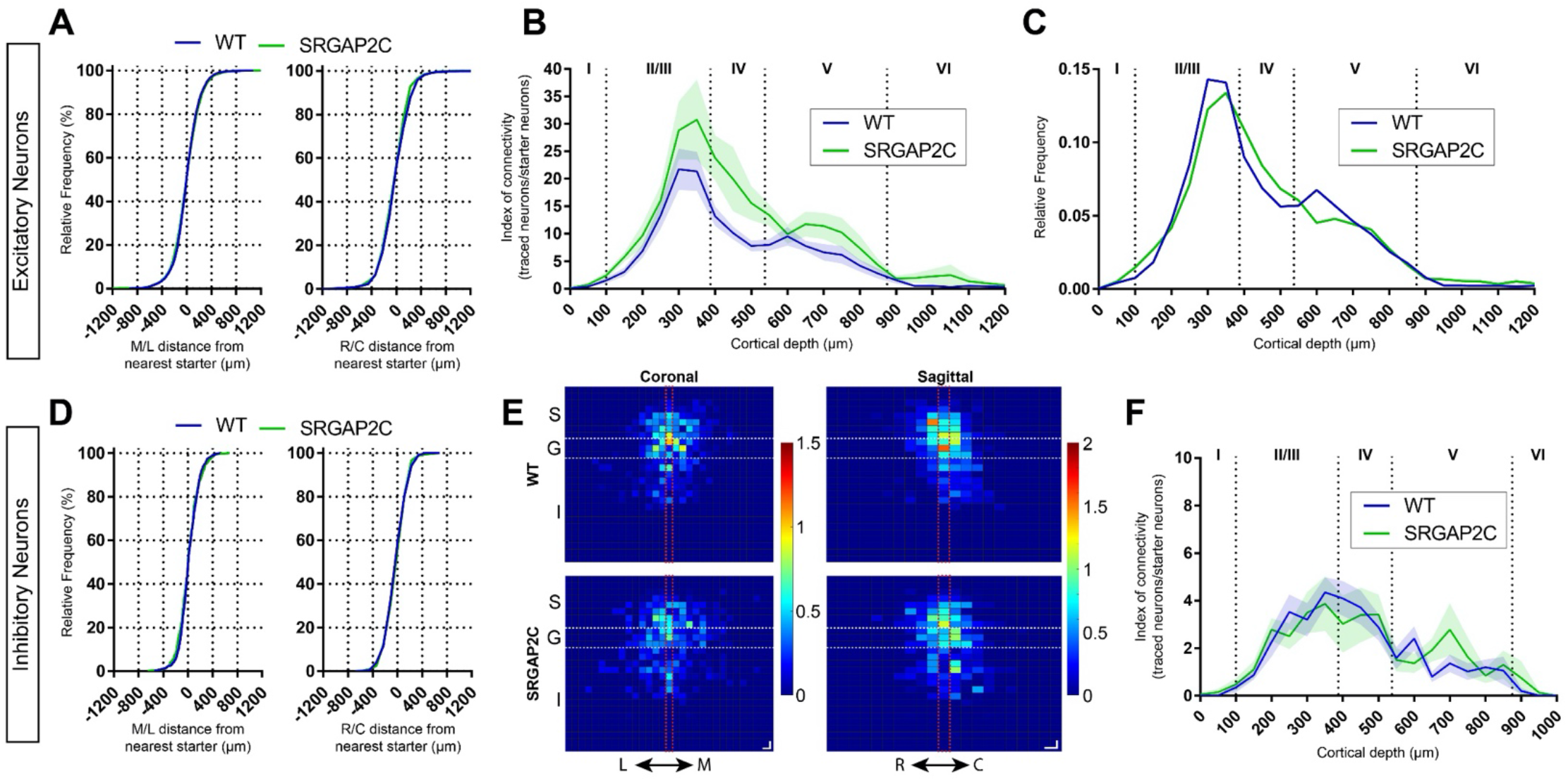
Distribution of RABV traced neurons locally in S1. (**A**) Distance between RABV traced excitatory neurons in S1 and their closest starter neuron along the medial/lateral (M/L) or rostral/caudal (R/C) plane. No difference was observed between WT and SRGAP2C mice. Data shown as relative frequency distribution. (**B**) Cortical layer distribution of RABV traced excitatory neurons in S1 shown as Index of connectivity (IOC, number of traced neurons / number of starter neurons). Shaded are indicates s.e.m. (**C**) Fraction of RABV traced neurons across cortical layers in S1. Dashed lines indicates borders between layers. Roman numbers identify cortical layers. (**D**) Same as (A), for inhibitory neurons. For analysis of interneurons, Parvalbumin-positive and Somatostatin-positive were grouped together. (**E**) Density plot showing distribution of traced inhibitory neurons relative to their closest starter neuron for coronal (left, L and M indicate lateral and medial orientation, respectively) and sagittal view (right, R and C indicate rostral and caudal orientation, respectively). Center bins aligned with relative position of starter neuron are indicated by red dashed lines. S, supragranular (layer 2/3), G, granular (layer 4), I, infragranular layers (layer 5/6). For coronal, bin size = 50×50 μm. For sagittal, bin size = 50×100 μm. Colors in density plots indicate IOC. (**F**) Same as in (B), for inhibitory neurons. Shaded are indicates s.e.m.

**Fig. S6.**
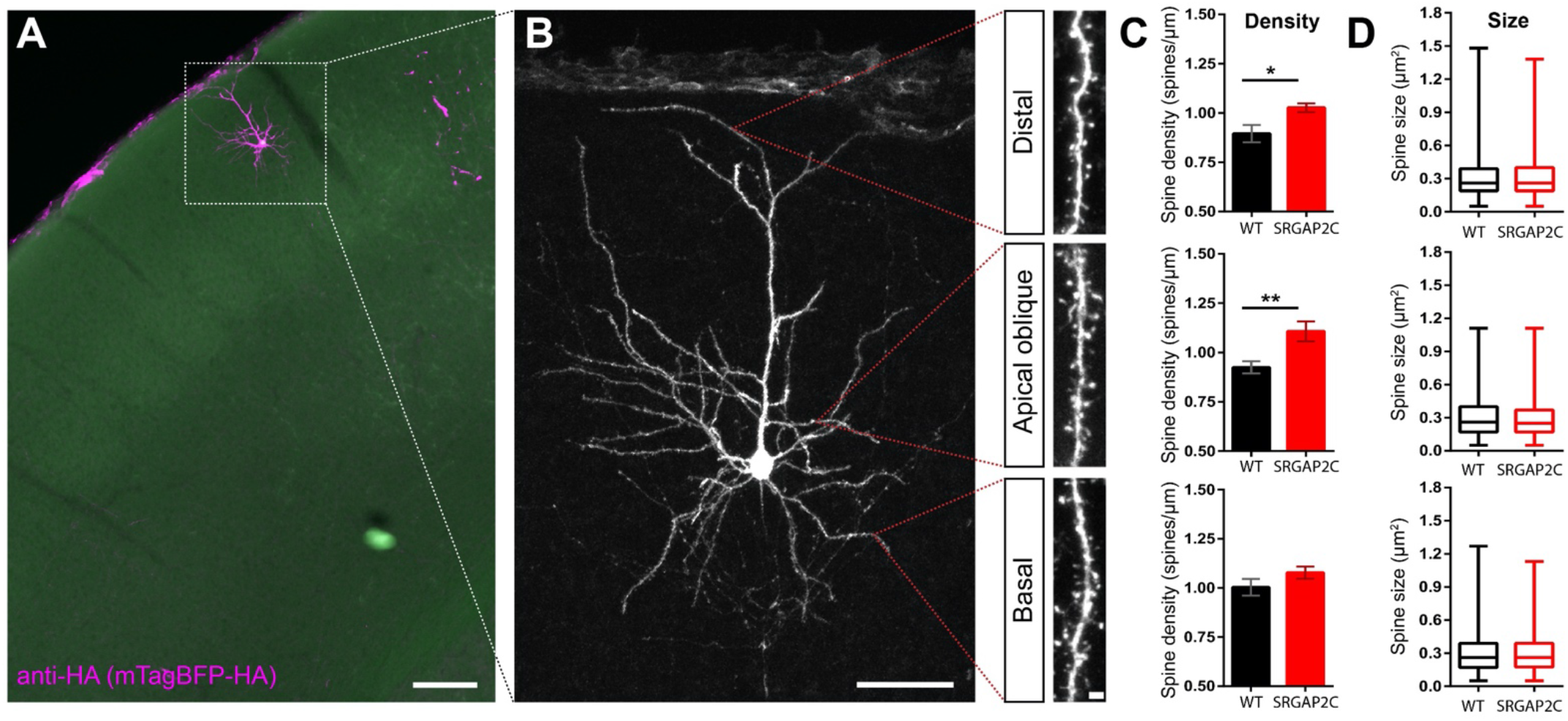
SRGAP2C expression selectively increases synaptic density on apical dendrites. (**A**) Coronal section stained for HA showing sparse labeling of a layer 2/3 cortical pyramidal neuron in the barrel field of the primary somatosensory cortex. Scale bar, 150 μm. (**B**) Higher magnification of neuron in (**A**). Red dotted lines indicate approximate location where spine density and size were quantified for distal, apical oblique, and basal dendritic compartments. Panels on right show high magnification images of dendritic segments on which spines can clearly be identified. Left panel scale bar, 50 μm. Right panel scale bar, 2 μm. (**C**) Spine density is increased for apical but not basal dendritic segments. (distal: *n* = 21 segments for WT and SRGAP2C, apical oblique: *n* = 33 segments for WT and *n* = 24 segments for SRGAP2C, basal: *n* = 32 segments for WT and n = 24 segments for SRGAP2C). Bar graph plotted as mean ± s.e.m. \**P* < 0.05, \*\**P* < 0.01. (**D**) Spine size is not significantly changed in adult SRGAP2C expressing layer 2/3 cortical pyramidal neurons. (distal: *n* = 1273 spines for WT and *n* = 1083 spines for SRGAP2C, apical oblique: *n* = 2401 spines for WT and *n* = 1650 spines for SRGAP2C, basal: *n* = 2286 spines for WT and *n* = 1448 spines for SRGAP2C). Bar graph plotted as mean ± s.e.m.

**Fig. S7.**
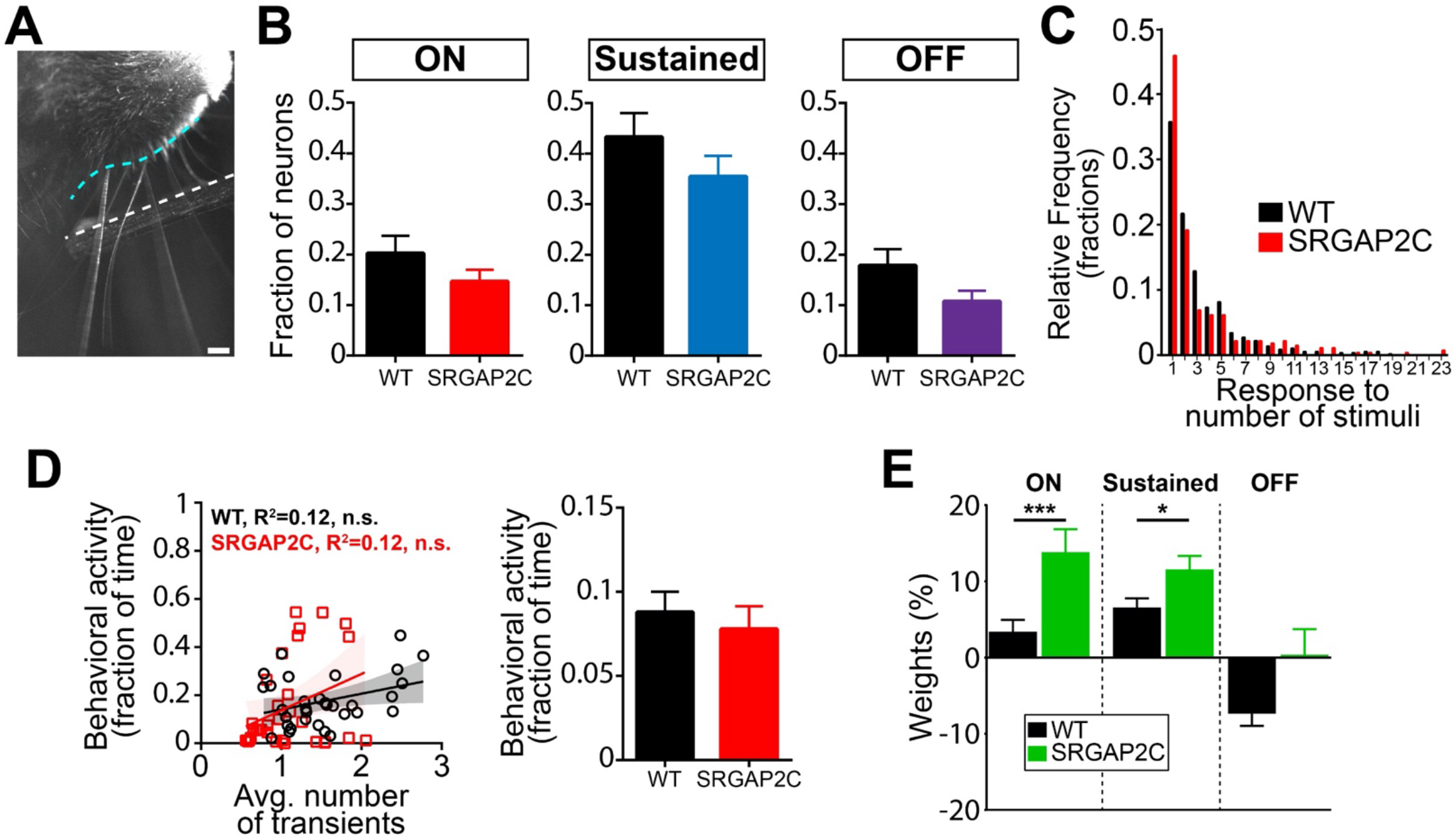
Neuronal activity in response to whisker stimuli. (**A**) Top-down view of placement of stimulating rod (white dashed line, 2mm away from the whisker pad) next to right whisker pad (cyan dashed line). Scale bar, 1mm. (**B**) Fraction of neurons responding to onset (ON), sustained phase, or offset (OFF) of whisker stimulus. (**C**) Frequency distribution of response fraction for neurons responding to either onset, sustained phase, or offset of the stimulus. (**D**) Left panel: fraction of time during which behavioral activity was observed. Right panel: correlation between behavioral activity and average number of transients. (**E**) Weights from the SVM classifier grouped by ON, Sustained, and OFF responses. Bar graph plotted as mean ± s.e.m. *\*P* < 0.05, *\*\*\*\*P* < 0.0001.

**Fig. S8.**
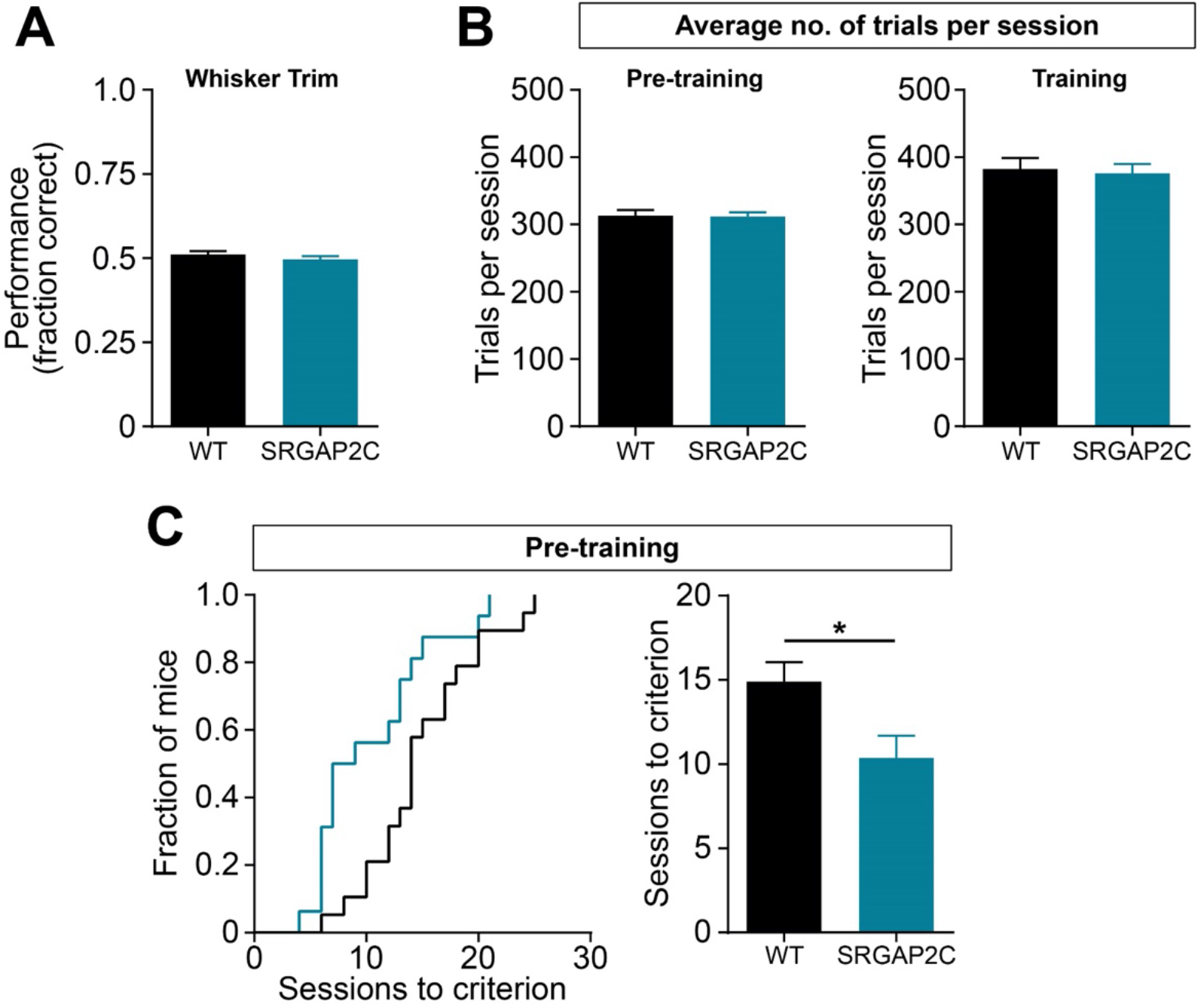
Whisker-based texture discrimination task. (**A**) Performance (fraction correct) after whiskers facing texture were trimmed was nearly 50% (chance level), showing that mice needed their whiskers to perform this task. (**B**) Average number of trials per session for pre-training and training phase. (**C**) Left: cumulative histogram for number of sessions to learning criterion during pre-training. Right: Average number of sessions to reach learning criterion during pre-training. Bar graph plotted as mean ± s.e.m. \**P* < 0.05.

**Movie S1. 3D reconstruction of a representative RABV traced brain mapped onto Allen Reference Atlas.**

Movie of a representative reconstructed RABV traced brain. Reference brain was adapted from the Allen Institute. Octahedrons represent RABV traced neurons and are color coded based on their anatomical location. See text for details.

**Movie S2. Whisker based texture-discrimination task**

Representative movie of a head-fixed mouse performing in a whisker-based texture discrimination task. Textures are rotated into position and subsequently moved towards the whisker pad of the mouse. A water reward is received when the mouse responds by licking the correct left or right lick port. Incorrect responses are punished with a timeout. The task is performed in the dark and infrared illumination was used for video recording.

